# Global StationaryOT: Trajectory inference for aging time courses of single-cell snapshots

**DOI:** 10.64898/2025.12.18.694987

**Authors:** Cole Boyle, Elias Ventre, Geoffrey Schiebinger

## Abstract

Trajectory inference (TI) methods for single-cell snapshots of developmental systems have yielded numerous insights into the gene regulatory networks (GRNs) that control cell differentiation. Many TI algorithms have been proposed for recovering cell trajectories from single samples containing cells spanning a spectrum of differentiation states; however, these methods cannot leverage temporal information when a time course of such diverse samples is available. As interest grows in understanding how the regulation of GRNs changes as an organism ages, current TI theory and methods must be adapted to take advantage of all information in aging time courses of single-cell data. In this paper, we present our novel age-conscious method, global StationaryOT, which exploits the temporal information in aging time courses to simultaneously reconstruct debiased cell trajectories at all ages. We demonstrate that this first-of-its-kind method achieves more accurate, biologically consistent trajectories in synthetic and real biological contexts where data sparsity produces significant noise in the outputs of current TI methods when they are applied to time course samples independently.

## 1 Introduction

Trajectory inference (TI) is a staple method for predicting the genes driving cell differentiation from single-cell data sets. Numerous methods have been proposed across a wide array of application domains [1–6], many of which have led to new discoveries by identifying or filling in the gene regulatory networks (GRNs) responsible for driving cells towards committed types. However, since these methods operate on data sampled around a single point in time, questions often remain as to the manner in which the regulation of these GRNs changes as an organism ages. To answer these questions, experimentalists have begun producing time courses of developing cells from organisms at various ages [7]; see Figure 1 for a simulated example. TI analyses of these time courses have thus far treated each sample independently, running the risk of over-fitting sampling noise that can degrade age-related signals. In this work, we propose a novel TI method specifically designed to reconstruct cell trajectories at all ages in these time courses simultaneously. Our method shares information globally across all ages, smoothing disparities in the relative abundance of cell types caused by sampling effects and allowing for more consistent trajectory estimates and the recovery of clearer aging signals.

**Figure 1.**
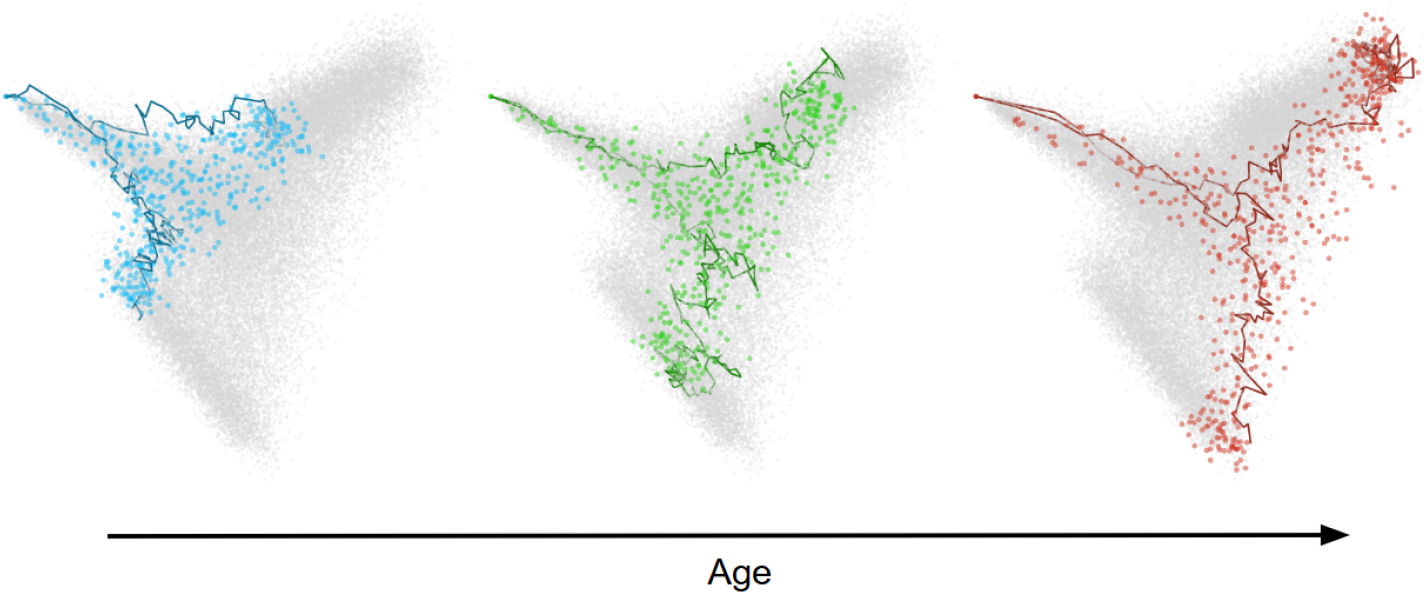
Simulated two-lineage aging single-cell time course. As the system ages, the lineages extend and rotate in the gene expression space. Sampled data at initial, intermediate, and final ages are coloured, along with sample cell trajectories, while data sampled at other ages are shown in grey.

At a given organism age, the persistent and stable cycle of cell growth, differentiation, and death allows many developmental processes to be viewed as near stationary; i.e., the population of cells across differentiation states is in equilibrium over short time scales. With this assumption, we model the developmental system at all ages as quasi-static [8, 9]. We reason that under normal homeostatic conditions, organisms rely on having steady populations of differentiated cells, so any age-related changes cannot occur quickly enough to significantly disrupt the balance of cell states at any age. This quasi-static assumption underlies our method’s approach to reconstruction and regularization.

At any age, we can recover cell trajectories via the mass displacement necessary to restore equilibrium after a short period of growth [4, 5]. We can then debias this reconstruction process using temporal information by (i) imposing that the system evolve slowly enough such that the cell distribution does not change rapidly between consecutive ages and (ii) strictly enforcing the equilibrium condition at each age. The first form of regularization ensures a smooth transition between the reconstructions, reducing overfitting due to general sampling noise. The second form is critical in cases where progenitor and terminal cell states are missing from our samples at certain ages. The imposition of equilibrium can force the inclusion of these states in the reconstruction when they are absent from a particular age, but present in nearby samples. Our method, global StationaryOT, balances these regularization principles with a data-fitting term to provide debiased trajectory reconstructions at all ages across a time course.

### Related work

The first trajectory inference techniques were developed for single-cell snapshots of a developmental system sampled at a single age [1, 2]. Since cells in these snapshots are unordered, a critical goal for these methods was the (mostly) unsupervised inference of a so-called pseudotime that sorts cells by their state of differentiation. Later methods enabled improved ordering estimates by utilizing additional information, such as RNA velocity [3] or cell growth rates [5]. Modelling single snapshots as coming from a stationary system was originally proposed by Weinreb et al. and used to reconstruct cell paths via their method Population Balance Analysis (PBA) [4]. This assumption was later used by Zhang et al. [5] to build their optimal transport-based method StationaryOT, on which our method is based.

After the original TI methods were proposed for snapshot data, it was realized that inference could be further improved by sampling cells as they develop in real-time [10, 11]. Such real-time time courses are formed by periodically sampling a population of developing cells so that the samples are already approximately sorted by differentiation state, allowing TI to proceed by linking cells taken at consecutive times [10]. In contrast, we consider time courses of snapshots that ideally consist of cells across a spectrum of differentiation states, such that without regularization, cell trajectories are confined to the states present in a single sample. The setting we deal with can be viewed as an abstraction of the one considered when developing previous time course based methods, as we have access to time-stamped cells, but these time stamps are no longer (necessarily) correlated with the development stage of the cells. In particular, our method extends the regularization ideas developed for real-time time courses, particularly those of Lavenant et al. [12] and their Wasserstein regression scheme, to the setting of quasi-static aging. An illustration of the differences between these settings and methods is given in Figure SI.1.

## 2 Methods

### 2.1 Quasi-static modelling of aging cell development

Let *X* ⊂ ℝ^*G*^ be the space of all possible gene expression states that cells can take across *G* genes. Each vector component represents the expression level of a particular gene. The effects of aging are modelled as a slowly evolving perturbation of the continuous developmental system, in which cell differentiation occurs significantly faster than the accrual of noticeable aging effects. As such, we introduce two time scales: the organism’s chronological age *t*, and its biological or physiological age *a* = *ct*, where *c* is the rate at which the organism ages. A specific cell at time *t* is modelled as a point, *X*_*t*_ ∈ *X*, with a trajectory described by the following stochastic equation

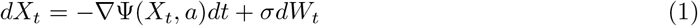

where Ψ is an unknown epigenetic landscape. We have expanded upon previous models [4, 5] by allowing Ψ to evolve slowly with age. This addition is consistent with the observed correlation between structural epigenetic changes and an organism’s age [13–17]. *σdW*_*t*_ is an increment of standard Brownian motion with diffusivity *σ*^2^, modelling the unaccounted for perturbations impacting the cell’s velocity, e.g. thermal fluctuations influencing the rate of DNA transcription. Cells also divide and exit the system with age-dependent exponential rates *α*(*X*_*t*_, *a*) and *β*(*X*_*t*_, *a*), respectively, i.e., over an infinitesimal interval, *dt*, they have a probability *α*(*X*_*t*_, *a*)*dt* of dividing and probability *β*(*X*_*t*_, *a*)*dt* of dying or leaving the system.

Given this model for individual cell trajectories with birth and death, the population distribution, *ρ*(*x, t*), over *X* of all cells at age *a* satisfies the following population balance PDE: ∂_*t*_*ρ*(*x, t*) = *Lρ*(*x, t*), with *L* given by

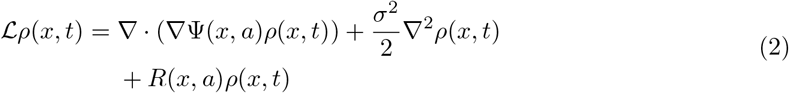

where *R*(*x, a*) = *α*(*x, a*) − *β*(*x, a*) is the age-dependent growth rate of cells at the gene-expression state *x* ∈ *X*. States with *R*(*x, a*) *>* 0 are sources, where cells are actively proliferating, while states with *R*(*x, a*) *<* 0 are sinks, as cells may die or leave the system in such locations.

Now, suppose the organism has reached a biological age *a* and has populations of cells spanning all levels of differentiation. Let the average lifespan of cells born at this age be denoted by *ℓ*(*a*). In developmental systems for which there is a continuous cycle of cell division, differentiation, and death, if the aging rate, *c*, is small enough, the system will remain in approximate equilibrium over the course of a single cell life cycle. More precisely, over a time interval, Δ*τ* = *O*(*ℓ*(*a*)), if *ℓ*(*a*) ≪ 1, then Δ*a* = *c*Δ*τ* ≪ 1. So *a*, and hence the system, will have undergone a negligible change over Δ*τ* and yet individual cells will, on average, have undergone a complete life cycle. Thus, over the interval (*t, t* + Δ*τ*), the normalized population distribution 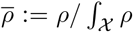 will remain close to a steady-state distribution, 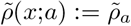, obtained from solving

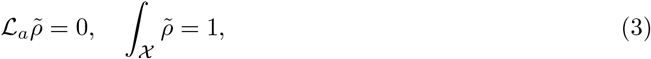

where *L*_*a*_ is obtained from *L* by treating *a* as a fixed parameter. In the thermodynamics literature, such a system is considered to be quasi-static [8, 9], or “slow” [18], particularly in the limit when the process (here aging) occurs infinitely slowly and the system is precisely described by an infinite sequence of steady state distributions, such as those solving (3). Since it is critical to our trajectory recovery method, we state this quasi-static assumption (QSA) explicitly as

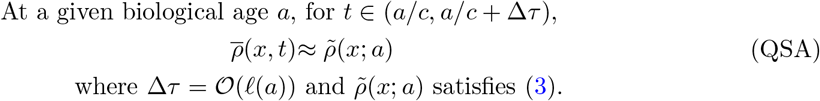

We note that the quasi-static assumption does not apply to systems that have been sampled during periods of rapid change or to those experiencing a sudden shock, e.g. due to sickness or serious injury.

Having described the system’s behaviour over short periods of time, we move to a characterization of the aging dynamics. Letting 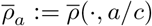, the curve defined by the function *a* 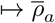 continuously conserves mass and hence will satisfy a continuity equation

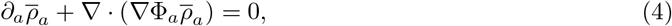

where Φ_*a*_(*x*) := Φ(*x, a*) is a potential guiding the aging quasi-steady-state distribution. Note that, unlike Ψ, the potential Φ is not steering the differentiation of individual cells, but rather the entire distribution of developing cells, including fluctuations in cell densities due to division and death. Now, by identifying the density 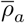 with its corresponding probability distribution on *X*, 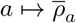 can be viewed as an absolutely continuous curve in the Wasserstein-2 space of probability distributions, *W*_2_(*X*) [19]. Thus, when Δ*a* is sufficiently small, 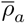 and 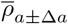 should be close with respect to the Wasserstein-2 distance, W_2_, and the movement of the distribution from 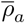 to 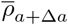 can be approximated using the optimal coupling.

#### 2.1.1 Observation Model

Given the above model, experimental observations of the system at an organism’s chronological age *t*, or equivalently, the biological age *a* = *ct*, are finite samples drawn from the marginal distribution 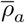 by sequencing cells. That is, if we measure the state of *n*_*a*_ cells 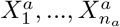 at age *a* then 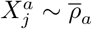. We can represent this sample as an empirical distribution, 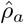 supported on 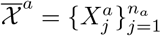

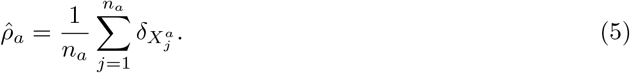

where *δ*_*x*_ represents the Dirac point mass located at state *x* ∈ *X*. Repeating the sequencing experiment on the organism at multiple ages, *a*_1_, …, *a*_*T*_, gives rise to a time series of such empirical distributions, 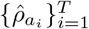.

In the following, we also assume knowledge of the cell growth rates *R*_*ij*_ := *R*(*X*_*j*_, *a*_*i*_), in the form of estimates denoted by 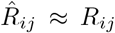. These growth rates can be estimated using cell cycle and apoptosis gene signatures to identify proliferating and dying cells [10]. Alternatively, prior biological knowledge of cell type specific growth can also be used once cell identities have been inferred with marker genes [5, 4, 20]. While our method can exploit age-dependent growth rates, currently estimates are typically only available at the age the cell was measured, i.e. we will need to take 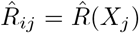 for all *i*. We adopt this approach in all our experiments below. Note that for brevity, we will often replace the subscript *a*_*i*_ with *i*, e.g. 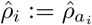.

### 2.2 Methodology Overview

Given a time course of single-cell samples, our goal is to recover estimates of the complete cell paths at every age we sample. Under the model proposed above, we will see below that this is possible provided we have estimates of the cell growth rates, 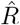, and noise level *σ*^2^. Our method builds off previous equilibrium based trajectory inference techniques designed for single stationary samples [4, 5]. Given cell growth rates, these methods recover cell trajectories as the necessary cell flow required to restore the population equilibrium after a short period of growth. Specifically, after cells divide, there will be more cell mass on high-growth regions, *R*(*x, a*) *>* 0, than low-growth regions, *R*(*x, a*) *<* 0. Absent cell movement, this growth would push the distribution out of equilibrium, or in our case, quasi-equilibrium. Hence, the cells’ flow can be recovered as the motion necessary to keep the population in a (quasi)-equilibrium. These methods fit a Markov transition matrix to the sampled data, 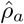, that approximates this flow and from which cell trajectories that trace paths from the high-growth to low-growth states can be sampled. Moreover, it can be used to compute the probability that any given cell in 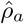 ends up in a particular low-growth region, e.g., that it leaves the system as a particular terminal cell type.

Critically, these reconstruction procedures rely on the samples, 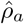, containing cells spread across the entire spectrum of differentiation states. When the 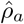 are sparse the inferred trajectories fail to correctly span this spectrum. In particular, missing cells in critical transition states will leave gaps in our trajectories, potentially causing spurious transitions. More importantly, when the samples fail to capture cells from every source and terminal region we may end up missing entire cell lineages (see the differences between Panels A and C in Figure 2 for an example) — if our sample contains cells of only a single type we cannot reconstruct the dynamics at all.

**Figure 2.**
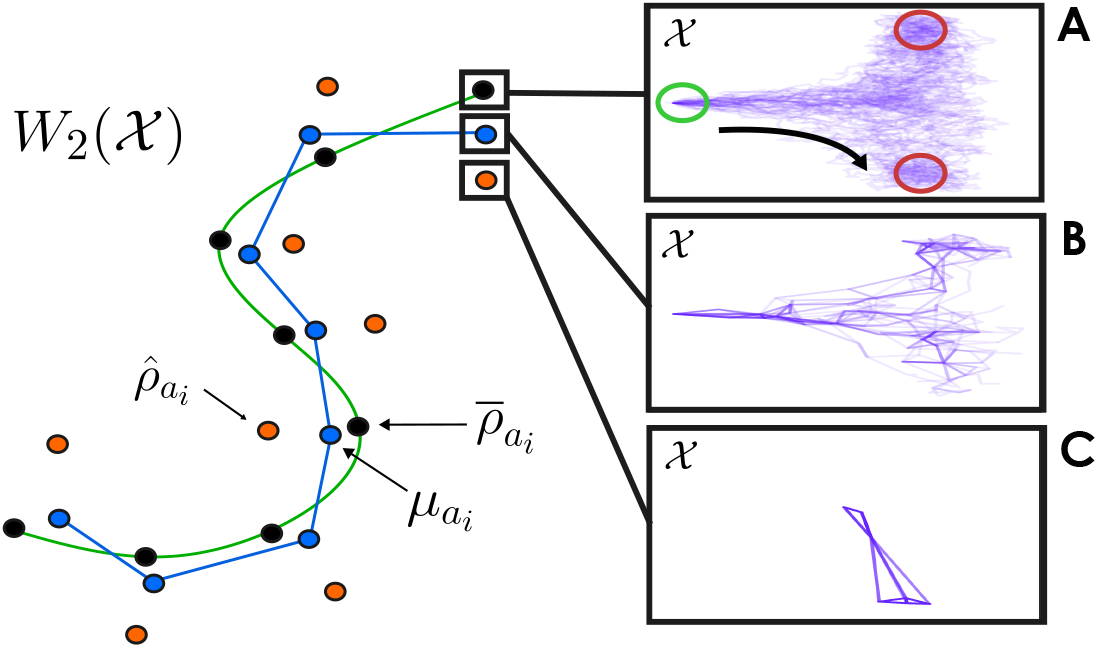
An abstract illustration of the quasi-static aging cell population curve, with sample and inferred cell paths displayed at the initial age. The green curve represents the aging cell population evolving according to (4) in the Wasserstein space, W_2_(X). At a particular age, a_i_, the true normalized cell distribution is represented by a black dot. Orange dots represent our finite sample distributions, 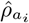, taken by sequencing cells, 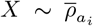. Since they are finite, they can be quite far from the true distribution 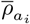. Blue dots represent the cell distributions, 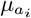, reconstructed at each age using global StationaryOT (gStatOT). These reconstructions are connected to highlight gStatOT’s approach for improving inference by sharing information across samples to (i) smooth transitions in the population distribution between ages and (ii) balance cell growth at each age. We implement (i) by minimizing the smoothing term, in (14), which can be seen as attempting to bring the blue points closer together to better approximate the green curve with the blue curve, an approach analogous to the smoothing procedures used in standard regression problems to address overfitting. With (ii) we aim to use our QSA to infer more robust cell paths at each age using the chain of approximations 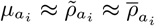, where 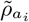 is the solution to (3). Panel **A** shows the true cell trajectories at age a_1_, traversed from primary sources (green region) to sink regions (red) in the gene space, X. gStatOT and StationaryOT (StatOT) trajectory reconstructions at the same age are shown in **B** and **C** respectively. The missing structure in **C** results from the single sample not containing enough information on its own (i.e. source and sink states) for StatOT to fully reconstruct both cell lineages starting from the primary source region.

When a time course of samples is available our method generalizes the equilibrium approach to trajectory inference to the quasi-static setting and aims to alleviate overfitting sparsity by sharing all available information when fitting the transition matrices at each age. We accomplish this by formulating a constrained convex optimization problem over (unnormalized) transition matrices, 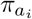, that balances data fitting and regularization to fill gaps in the data as well as pull the inferred dynamics in line with those of the ground truth. Specifically, we aim to fit each 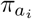 to all available data so that it best approximates the pure equilibrium dynamics specified in (3), and hence by our QSA also approximates the true dynamics at each age specified by the population model in (2). In particular, we reconstruct an optimal equilibrium distribution, 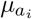, that better approximates 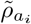 than our raw data 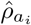, while simultaneously fitting the cell dynamics, 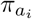, consistent with (3) and having solution 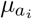. Consequently, we use the chain of approximations 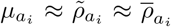 to build a curve *a* ↦ *µ*_*a*_, with associated equilibrium cell dynamics, *π*_*a*_, that approximate the true aging cell distribution curve 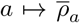 and the ground truth cell dynamics (2). See Figure 2 for a conceptual visualization of these curves.

### 2.3 StationaryOT

Since our method extends StationaryOT (StatOT) [5] to time series data, we briefly outline its methodology below. For a more detailed elaboration of the method, with benchmarks and the biological insights it has led to, see the texts [7, 5, 21].

Suppose that at age *a*, we know the stationary distribution 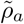, solving 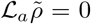. Although the population distribution remains unchanged, individual cells are still undergoing a cycle of division, differentiation, and death. Over a short interval Δ*t* < *ℓ*(*a*), the StatOT method observes that these cell dynamics can be approximated by separating the effects of cell flux and transport, after which we can recognize that the resulting transport equation for cell movement can be solved using entropic optimal transport (EOT).

Specifically, *L*_*a*_ is split into equations for growth and transport in the distributions 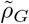 and 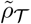 respectively. Setting *t*_0_ = *a/c*, these are given by,

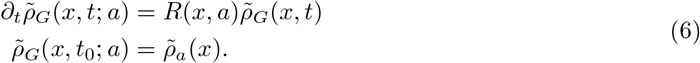

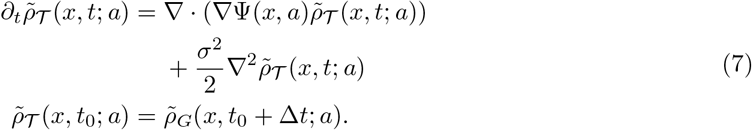

Notice that (6) applies the effects of cell growth to 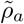, then, over this same period, Δ*t*, (7) specifies the transport of this evolved distribution via a drift-diffusion process. The solution to (7), evaluated at *t*_0_ + Δ*t*, is a *splitting approximation* of the equilibrium distribution 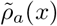. The two distributions coincide as Δ*t* tends to 0, with a pointwise error of order *O*((Δ*t*)^2^) [22], that is,

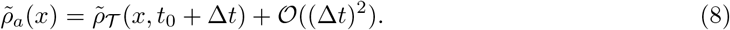

Proceeding to solve these equations, by denoting *g*(*x*; *a*, Δ*t*) = *e*^*R*(*x,a*)Δ*t*^, we can see that the solution to (6), evaluated at *t*_0_ + Δ*t*, is given by 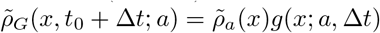.

With this solution, (7) and (8) specify that our cell dynamics over the period Δ*t* are approximated by a drift-diffusion process with initial distribution 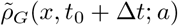 and final distribution 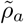. Note that on short time scales, the diffusion 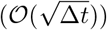 dominates drift (*O*(Δ*t*)) in (7). These diffusion-driven dynamics can be described by an EOT coupling, 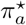, of 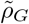 and 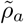. (This is called a Schrödinger bridge; see [23, 24] for a review of this connection between diffusion and EOT). More precisely, the coupling 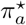 estimating cell dynamics is the unique minimum of the convex optimization problem,

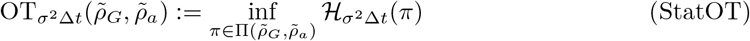

where Π(*γ*_1_, *γ*_2_) = {*π* ∈ *P*(*X* × *X*) : *P*_1#_*π* = *γ*_1_, *P*_2#_*π* = *γ*_2_} is the set of all couplings between the distributions *γ*_1_ and *γ*_2_. Here, *P*_*i*_ denotes the projection of a point *x* ∈ ℝ^*G*^ onto its *i*^th^ coordinate and _#_ denotes the push-forward operator so that *P*_*i*#_*π* = ∫*π*(*dx*_*i*_, ·) is the *i*^th^ marginal of *π*. Next,

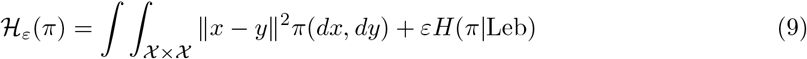

is the quadratic optimal transport loss with entropic regularization of strength *ε*. Leb is the Lebesgue measure on *X*^2^ and 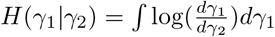 is the relative entropy between two distributions. To summarize the StatOT recovery process for a discrete sample of observations, 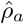, if we have growth rate estimates 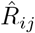 for the observed cells 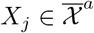 and an estimate of Δ*t* and noise level *σ*^2^ (see SI.2 for details on choosing these parameters), we first form the distribution 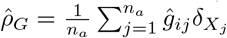 where 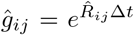. We then couple 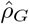 and 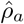 with entropic optimal transport using the regularization parameter *ε* = *σ*^2^Δ*t* to obtain an estimate for the cell dynamics in the form of a coupling matrix 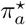. After row-normalization, we can sample cell trajectories and compute cell-fate probabilities. See SI.3 for a description of how we compute these probabilities using the cell growth rates.

### 2.4 Global StationaryOT

As discussed in the overview, given measurements of the developmental process at a series of ages, 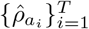, we propose a convex optimization problem that balances data fitting and regularization to reconstruct improved estimates of the 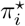 by smoothing their supports, namely the equilibrium distribution 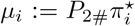, across all the sampled ages. Letting 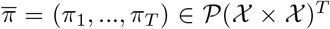 be a vector of couplings abbreviating *g*_*i*_ := *g*(·; *a*_*i*_, Δ*t*), our recovery program has the following form

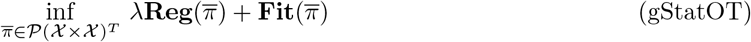

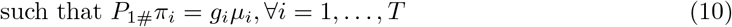

where the **Fit** operator is an abstraction of the StatOT problem, allowing the second marginal of each *π*_*i*_ to deviate from 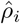, while **Reg** encourages smooth transitions between the support of the *π*_*i*_, consistent with a slow aging process.

#### 2.4.1 Constraint enforces equilibrium dynamics

Since the marginals of the *π*_*i*_ can now move during optimization, the constraint ensures that the first marginal of *π*_*i*_, *P*_1#_*π*_*i*_, still represents an evolution of *µ*_*i*_ after a period of growth, i.e. we still enforce that *P*_1#_*π*_*i*_ be a solution to (6) (evaluated at *a/c*+Δ*t*). Moreover, together with the unity constraint, *π*_*i*_ ∈ *P*(*X* × *X*), (10) implies that our reconstructed *µ*_*i*_ will be equilibrium distributions for the given growth rates. Namely, each *µ*_*i*_ must specify balanced mass on source and sink regions such that the net cell flux is 0. Indeed, both constraints together imply that we must have ∫*P*_1#_*π*_*i*_ = ∫*g*_*i*_*µ*_*i*_ = 1 ∫ *µ*_*i*_ = 1. So splitting the domain as *X* = *X*_source_ ∪ *X*_sink_ ∪ *X*_0_ for regions with *R*_*i*_ *>* 0, *R*_*i*_ *<* 0 and *R*_*i*_ = 0, respectively, and noting that *g*_*i*_ = 1 on *X*_0_, we can see that

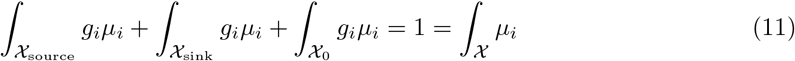

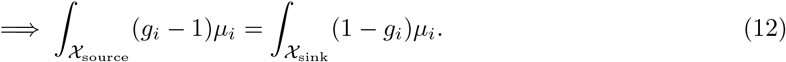

The left-hand side is the mass gained after growth, while the right-hand side represents what is lost. We are thereby regularizing the *µ*_*i*_, and thus *π*_*i*_, by strongly enforcing our equilibrium hypothesis that cells leaving the system are fully replaced at each age. As noted in the overview, this is critical, as equilibrium-based inference fails when a sample lacks source or sink states. In such cases, our *µ*_*i*_ will adapt and become equilibrium distributions by adding states from the missing regions in order to have ∫ *g*_*i*_*µ*_*i*_ reach unity. In practice, these additional states can come from other samples in our time course. Through this sharing of state information we are also able to satisfy the natural solvability criterion that arises from (12) due to needing both sides of the equation to be nonzero (see SI.1.2 for a discussion of this criterion).

#### 2.4.2 Fitting functional optimizes dynamics at each age

We have constrained the *π*_*i*_ to describe equilibrium dynamics without a priori knowledge of what their marginals should be. Now, we move to describing how we fit these *π*_*i*_ to our time course so that they best approximate the the dynamics described by (3), and hence by our QSA, the true cell dynamics implied by (2).

As laid out in the StationaryOT section, for a given estimate of the true equilibrium distribution (in that case 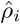 and in ours *µ*_*i*_) the cell dynamics consistent with (3) are obtained by minimizing the loss 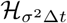 over couplings that, at least in theory, satisfy our constraint. To anchor these dynamics to each age, we keep each *µ*_*i*_ close to our sampled data 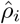, giving the data-fitting functional

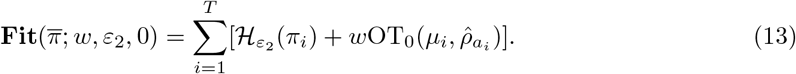

Here, each term in **Fit** extends the StatOT problem by penalizing the *µ*_*i*_ for deviating from 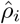 in squared W_2_ distance, instead of enforcing them to be equal. The strength of this penalty term is controlled by the parameter *w* such that, when *λ* = 0, taking *w* → ∞ recovers the StatOT problem, with the additional constraint of imposing balanced growth. In practice, we have found that simply taking *w* = 1 works well for sparse data, and so we set this as the default value for most of our experiments. Moreover, *ε*_2_ plays the same role as *ε* in StatOT and is set according to the same principles. See SI.2 for details on choosing these parameters, along with the smoothing parameter *λ*.

#### 2.4.3 Regularization functional smooths transitions between ages

Now, following our observation that in the quasi-static limit, consecutive 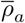 should be close with respect to the W_2_ metric, our QSA implies that the 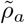 should be as well. Thus, to guide our reconstructed marginals, *µ*_*i*_, closer to 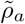 and hence 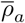 (as depicted in Figure 2) we define our **Reg** functional by the following

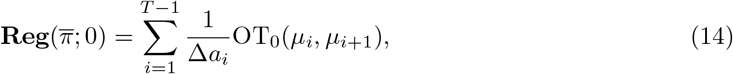

where Δ*a*_*i*_ = *a*_*i*+1_ − *a*_*i*_. While the evolution of the *µ*_*i*_ should happen on the aging timescale, since the rate *c* is generally unknown we absorb it into the regularization strength *λ*. So, in practice our regularization coefficient is given by 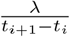, where *t*_*i*_ is the chronological age at which we sampled 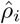. We emphasize that *t*_*i*+1_ − *t*_*i*_ is assumed to be significantly larger than our previously defined cell state transition time step Δ*t*.

#### 2.4.4 Implementation

To obtain a finite-dimensional convex optimization problem, we restrict gStatOT to measures supported on a finite set. Following Lavenant et al. [12], we choose this set to be the union of all our sampled data, 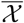, since a standard gridding approximation is intractable due to the high dimensionality of *X* (see SI.1.1 for details). Upon discretization, the gStatOT problem is strictly convex over the couplings, *π*_*i*_, and hence has a unique solution that we can solve for by maximizing the corresponding dual problem. We solve this dual problem using a block coordinate ascent (BCA) algorithm [25] with log-domain arithmetic, and use a tolerance on the duality gap as a stopping condition. See SI.1.3 and 1.4 for details on deriving the dual problem and our BCA implementation.

We provide Python implementations of gStatOT and StatOT using the JAX library [26–28], enabling seamless execution on GPUs or CPUs. Both implementations are designed to run on single-cell data stored in a standard AnnData [29] object and are available, along with detailed usage instructions on GitHub, at https://github.com/ColeBoyle/global-stationaryOT.

## 3 Results

### 3.1 Synthetic data: Aging epigenetic landscape

We demonstrate gStatOT’s ability to reconstruct trajectories in high-sampling-bias regimes on a simulated birth-death diffusion system representing the development and aging of two cell lineages in a 10-dimensional gene space. The differentiation of individual cells is driven by the dynamics in Equation 1 with an age-dependent developmental landscape, i.e. Ψ(*x, a*). This landscape generates a bifurcating developmental pathway from a single high-growth region towards one of two sink regions (as shown in Figure 2 Panel A). For clarity, we have centered the system so that the high-growth progenitors are located at the origin, and have chosen the unit of our chronological age, *t*, to be days. Cells are produced from this progenitor state and then develop by increasing their expression of Gene 1, followed by stochastically committing to one of two lineages by up- or down-regulating Gene 2, after which they eventually die. We refer to the two cell lineages as +2 and −2, respectively.

The developmental landscape’s dependence on age impacts the system by increasing the Gene 1 expression needed for commitment to occur, while also inducing an increase and corresponding decrease of Gene 3 in the +2 and −2 lineages, respectively. The precise mathematical details of our simulation can be found in SI.4.2. As in real data, only a minority of genes drive lineage development, while the rest add noise and dimensionality that confound inference. We simulated a time course with 25 ages by sampling all cells from the system every 4 days. Ground truth estimates of the cell trajectories and fate probabilities at each age were obtained by simulating trajectories starting from the sampled cell states at every age.

To introduce additional sampling bias, prior to applying our method we downsampled the time course to 10 cells per age. We applied gStatOT to the subsampled time course and compared the results to those obtained by applying StatOT to the same data at each individual age. See SI.4 for the details on how parameters were chosen and Figure SI.2 for an analysis of gStatOT’s sensitivity to errors in growth rate estimates. Figure 3 shows the subsampled time course and 25 reconstructed trajectories from both gStatOT and StatOT alongside 25 of the simulated trajectories at several ages. We observe that the reconstructed gStatOT trajectories more closely match those of the ground truth. Moreover, the growth rate regularization motivates the inclusion of the source cells located at the origin in gStatOT’s marginals, allowing us to correctly reconstruct trajectories that start from the origin. In contrast, StatOT’s trajectories are forced to start from positive-growth cells in the +2 lineage at age 30.

**Figure 3:**
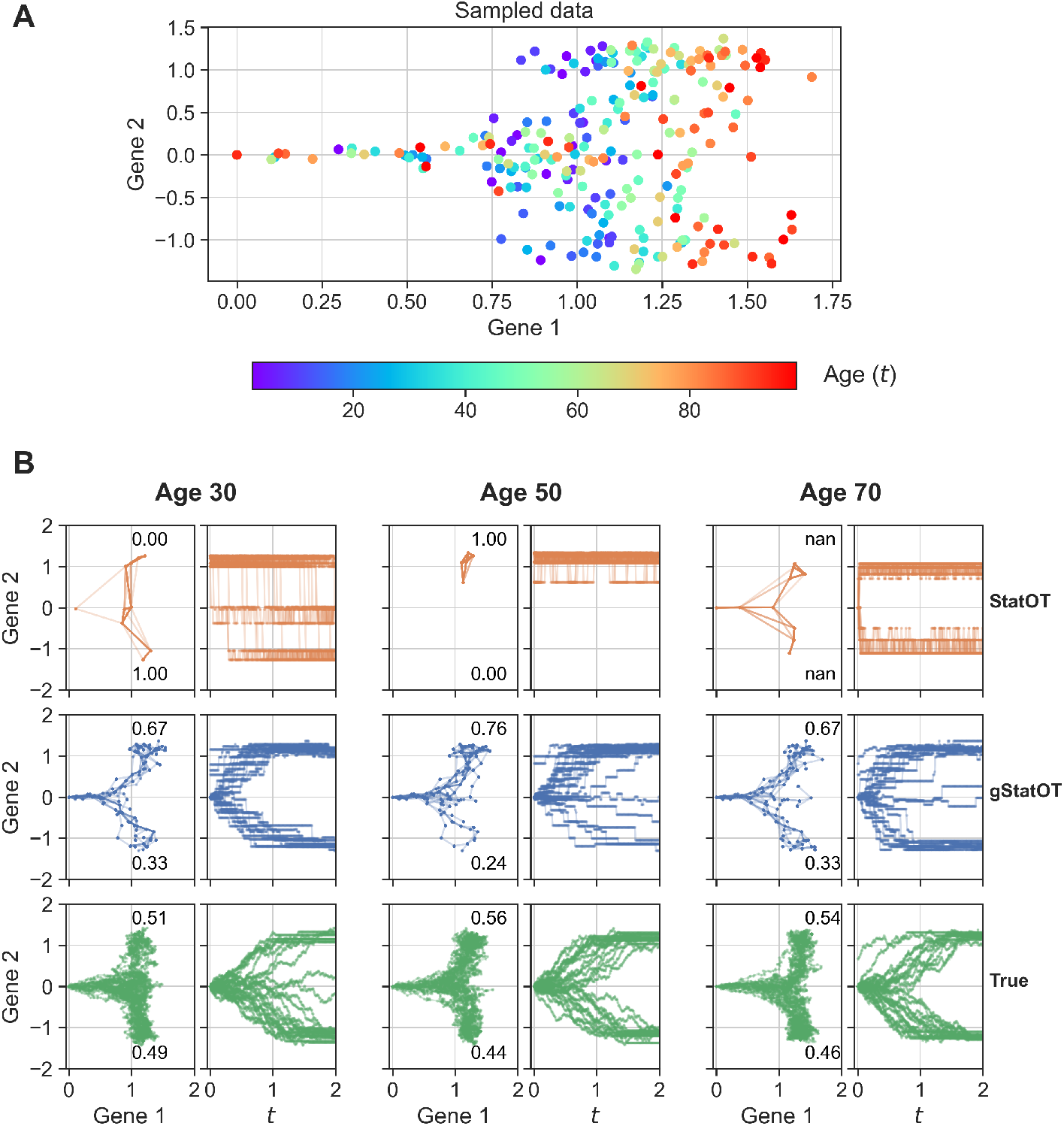
**A:** Subsampled simulated data at 25 ages with 10 cells per age. **B:** 25 2-day trajectories reconstructed from the top data using gStatOT and StatOT alongside 25 ground truth simulated trajectories at ages 30, 50, and 70. The adjacent plots show the (centered) Gene 2 expression of each trajectory as a function of its Gene 1 expression and chronological age, t, respectively. The numbers next to each lineage denote the proportion of trajectories that first reach either a +2 or −2 sink state. Note that there are no sink states in one, or both, of the lineages at each age. So while the StatOT trajectories can explore the state space at these ages, they are not able to reconstruct the full bifurcation structure of the system, and so, as with age 70, may fail to converge at all.

Next, we sought to quantify gStatOT’s performance and how well it improves with increased sampling rates. We used three metrics to evaluate the performance of our method, and compared the results to those obtained with StatOT and PBA [4]. With the first metric, we measure gStatOT’s ability to debias sparse samples by evaluating the W_2_ distance between the reconstructed marginals (*µ*_*i*_) and all simulated data from each age. StatOT and PBA operate exclusively on the sparse samples at each age, so to test if the *µ*_*i*_ are actual improvements over the raw data, we also show the W_2_ distance between these sparse samples and the full simulated data. The second metric compares the reconstructed trajectories to the ground truth cell paths we simulated at each age. As done in [5, 12], we choose to use the W_2_ metric on path distributions. Our final performance metric compares the inferred cell-fate probabilities with those obtained from the simulation in total variation (TV) distance. When a method failed to reconstruct fate probabilities we set the TV value for that age to its maximum value of 1.

Each model’s hyperparameters were chosen by optimizing the average of the three metrics on a single simulation (see SI.7.1). With these parameters, each method was then validated across 10 independent reseeded simulations, with the results being reported in Figure 4. A baseline was established for the marginal and trajectory metrics by comparing the complete simulated data from the initial simulation to the 10 reseeded ones. Panel **A** of Figure 4 shows that across all 25 sampled days in the time course, gStatOT was able to construct marginals closer to our ground truth simulated data than the subsamples and outperform StatOT and PBA in both the trajectory reconstruction and fate probability estimation. The results are summarized by the mean and standard deviation of each metric across the 10 independent simulations.

**Figure 4:**
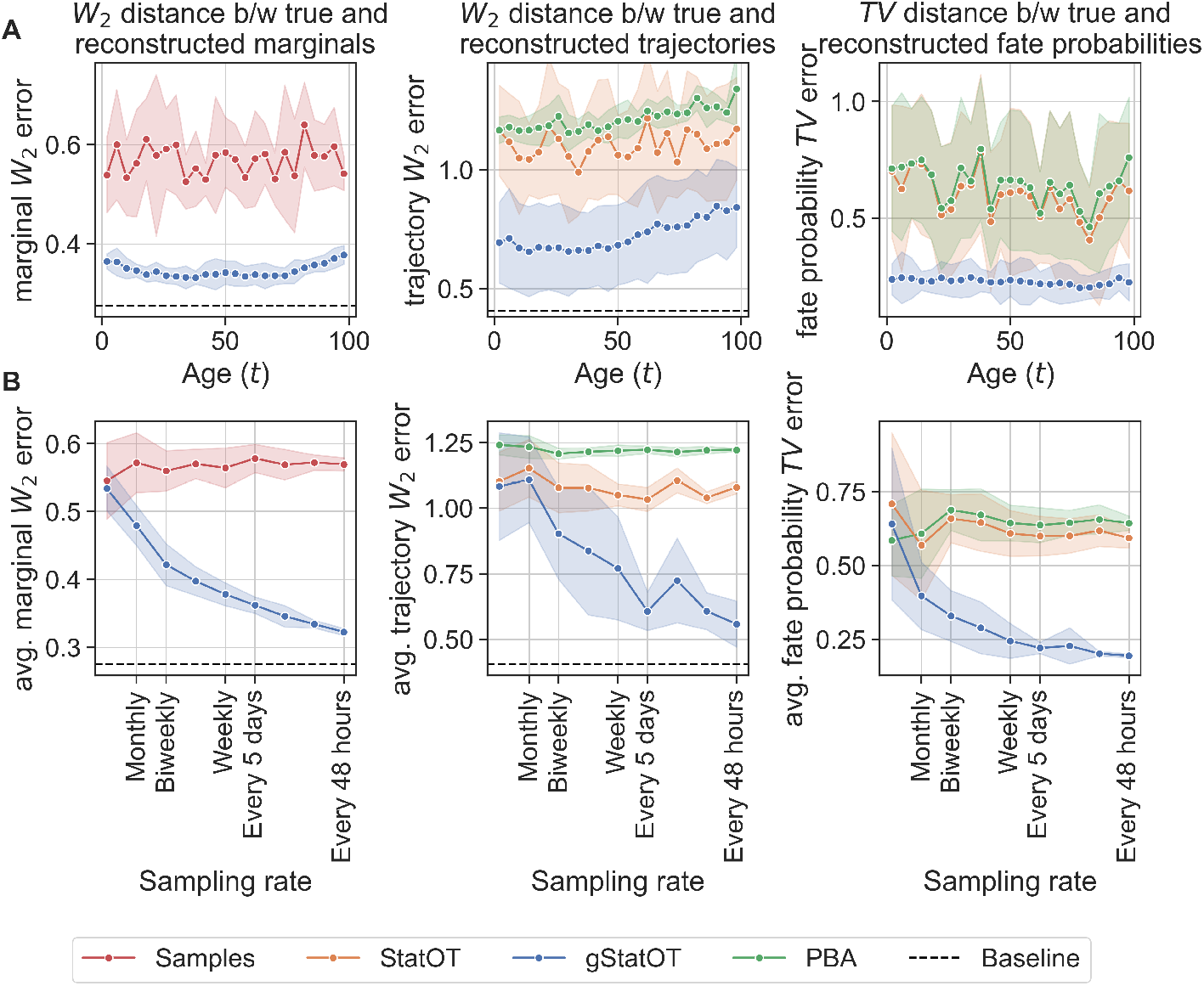
**A:** Model performance across age in marginal, trajectory, and fate probability reconstruction for simulated data with 25 ages and n_a_ = 10 cells per age. **B:** Model performance for varying number of intermediate time points. The decreasing trends in all plots demonstrate that gStatOT is successfully transmitting information across ages to improve its estimates. The data for each sampling rate is the average of each metric across all ages. The results are summarized by the mean and standard deviation of each method’s performance on 10 independent reseeded simulations. The baseline represents the average distance between the original simulated distributions and those obtained from the 10 reseeded simulations.

To test whether gStatOT is successfully exploiting information across multiple time points, we repeated the above process for time courses generated with variable sampling rates. Panel **B** of Figure 4 shows the average performance of gStatOT, StatOT and PBA across all ages for each time course. We observe that gStatOT’s estimates increase in accuracy for each metric as the sampling frequency increases and reach near baseline once we are sampling every other day. On the other hand, those of StatOT and PBA remain relatively constant, providing evidence that gStatOT is successfully utilizing global information across the time course.

### 3.2 Synthetic data: Loss of cellular identity over age induced by epigenetic erosion

Within several biological systems, a progressive loss of cell identity has been observed to occur over the course of the aging process [16, 30, 31]. Due to the effects this has on a cell’s ability to perform its specialized roles, it has been theorized that this loss of identity could play a critical role in bringing about the physiological changes and diseases generally associated with aging [17, 32]. A proposed mechanism for this loss of identity is the erosion of epigenetic structures that regulate gene expression. Structural elements such as chromatin normally help cells specialize by inhibiting or promoting gene expression depending on their placement on the DNA strand; however, they also play a role in DNA damage repair. As the need for repairing damage increases over time, chromatin loses its ability to regulate gene interactions, resulting in a loss of cell identity [17, 33].

To benchmark gStatOT on data from such a setting with an accessible ground truth, we developed a realistic simulation of a developmental system subject to progressive cell identity loss. We considered a branching system that develops according to the GRN in Figure 5 Panel **A**, which specifies a chain of genes controlling both cell growth and differentiation. Cells start out in a progenitor state defined by expression of the proliferation gene, *G*_*g*_, and the gene *G*_1_. They then progress into one of two lineages: branch 1 (B1) or branch 2 (B2), depending on which one of the mutually repressive genes *G*_*B*1_ or *G*_*B*2_ they express. We biased cells towards B1 by specifying a larger interaction strength between *G*_4_ and *G*_*B*1_. Upon terminally differentiating, the cells begin to express *G*_*d*_, which triggers apoptosis. To model the loss of cell identity with age, we made the repression strength, *S*_*B*1↔*B*2_, between the lineage driver genes *G*_*B*1_ and *G*_*B*2_, decrease linearly with age. This can be viewed as cells in intermediate states inheriting deteriorated epigenetic structures that fail to inhibit one or the other gene [34]. As a result cells are increasingly likely to terminate in an ambiguous state (denoted as B1+B2) by expressing both lineage markers *G*_*B*12_ and *G*_*B*22_.

**Figure 5.**
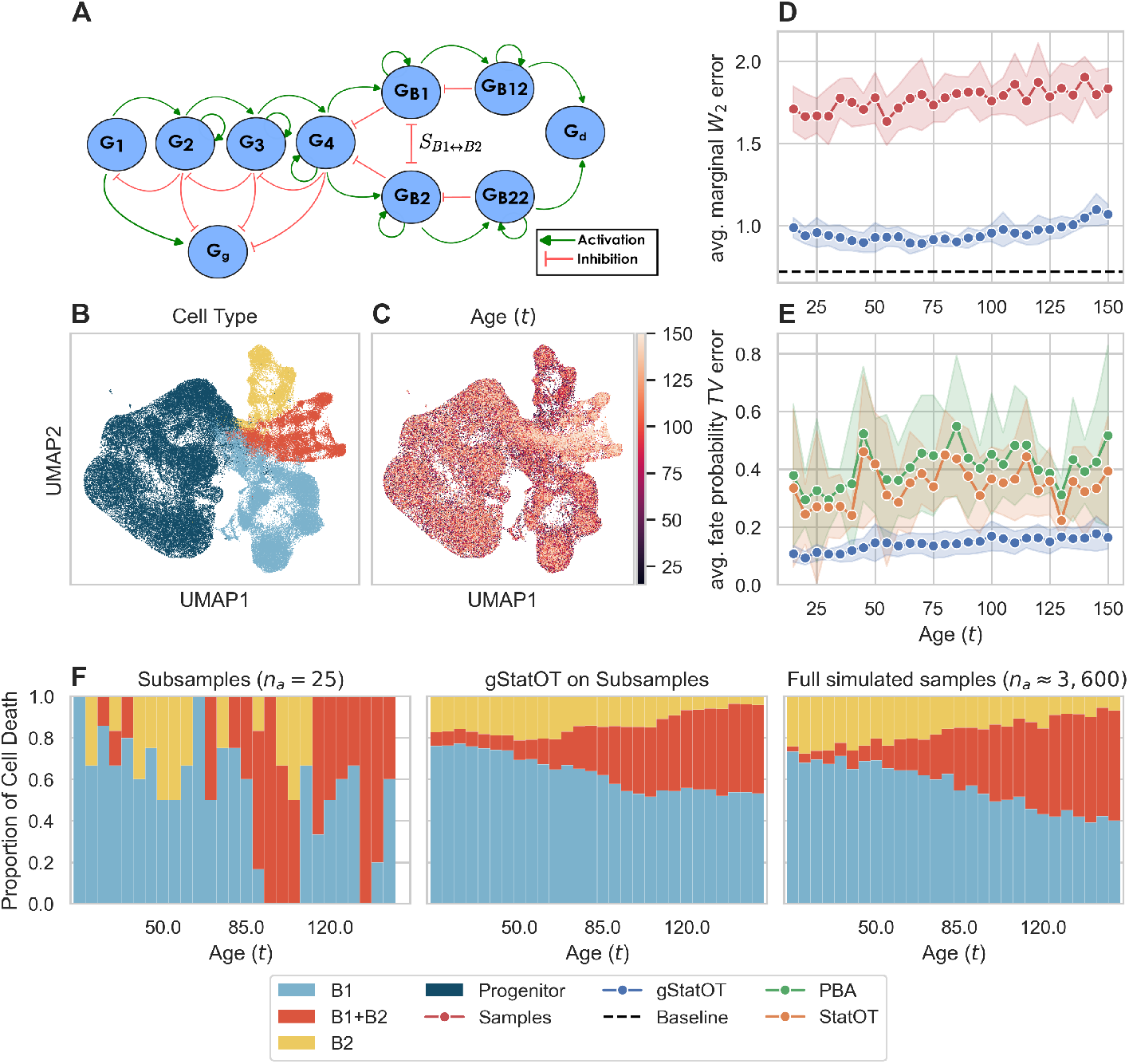
Simulated gene regulatory network (GRN) model for cell differentiation with progressive cell identity loss. **A:** The GRN we used to simulate the loss of cell identity with age. Cells start out in a progenitor state defined by expression of G_1_ and the proliferation gene G_g_. They then progress into one of two lineages, B1 or B2, depending on which of the mutually repressive genes G_B1_ or G_B2_ they express. S_B1↔B2_ represents the age-dependent strength of this repression. Upon terminally differentiating by expressing either of the lineage markers G_B12_ or G_B22_, cells begin to express G_d_, which triggers apoptosis. **B, C:** UMAP embedding of the simulated time course coloured by cell type and age, respectively. Note that the ambiguous B1+B2 branch consists of cells primarily sampled at later ages. **D:** Marginal distribution reconstruction performance for gStatOT on the downsampled data. **E:** Fate probability reconstruction performance for gStatOT, StatOT, and PBA on the downsampled data. Results are summarized by the mean and standard deviation of each metric across 10 reseeded simulations. **F:** Proportion of cells dying in each branch at each age as predicted by the sparse samples, gStatOT’s marginals, and the complete simulated data. See Figure SI.3 for a quantitative comparison of these proportions across all ages.

To realistically generate single-cell data from this system we employed the GRN simulator BoolODE that was developed as part of the BEELINE framework [35–37]. Since BoolODE does not simulate cell growth or allow for the regulatory strengths between genes to depend on time, we modified the stochastic ODE model it generated for our GRN by making the parameter *S*_*B*1↔*B*2_ depend on age, and then proceeded to simulate it with birth and death, similar to what was done in our previous example. We periodically sampled from the simulation to create a time course with 28 ages. Ground truth fate probabilities for all sampled cells were obtained using the same method as in the previous example. To further emulate the noise in real sequencing data, we also followed BoolODE’s protocol for giving each sample in the time course a dropout rate of 50%. See SI.4.3 for the full details of our simulation procedure.

Panels **B** and **C** in Figure 5 show the time course embedded in UMAP space [38], coloured by cell type (e.g. branch B1, B2, or B1+B2) and the age at which the sample was taken. Each cell type label was assigned based on each cell’s expression of the marker genes *G*_*B*12_ and/or *G*_*B*22_, after dropouts were added. As intended by our model, there is a mix of cells taken at all ages in branches B1 and B2, while the ambiguous B1+B2 branch consists of cells primarily sampled at later ages.

We downsampled the simulated data to 25 cells per age before applying gStatOT to test its marginal distribution and fate probability reconstruction ability. The results of these tests are shown in Panel **D** and Panel **E** of Figure 5. Parameters for each of the methods tested were fit to optimize fate probability reconstruction (see SI.4.3 for application details and SI.7.2 for the parameter sweeps). We can see that gStatOT’s marginal distributions are again closer to the full simulated data at each age than the sparse samples — approaching the baseline distance between reseeded simulations. In Panel **E** we see that gStatOT improves fate probability reconstruction across all ages when compared with StatOT and PBA. In particular, it significantly reduces variance in the TV error across repeated runs compared to that of StatOT and PBA, with these latter methods suffering from the limited availability of sink states (i.e. cell states that express the lineage markers *G*_*B*12_ and/or *G*_*B*22_ and have negative growth rate due to their expression of *G*_*d*_).

A property of potential biological interest in such systems is the proportion of cells dying in each state over the course of the aging process. Panel **F** illustrates these proportions, which were computed using the cell growth rates and each of the sparse samples, gStatOT’s marginals, and the complete simulated data. For example, with gStatOT, we compute the predicted number of cells dying in a branch *b* at age *a* as 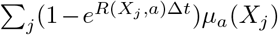, where the sum is taken over sink cells, *X*_*j*_, labeled as being a part of branch *b*. Here, we can see evidence that gStatOT’s constraint, enforcing mass balancing, is accurately correcting the sampling bias present in the sparse samples by placing mass on sink states so that the predicted proportion of cells dying in each of the branches better matches that of the ground truth. See Figure SI.3 for a quantitative comparison of these proportions across all ages.

### 3.3 Human Hematopoiesis

For an initial test of our method on real biological data, we applied gStatOT to a hematopoietic scRNA-seq time course that spans the entire human lifetime [7]. The time course consists of 22 time points: 8 gestational, taken between 10 and 23 weeks post-conception; 13 postnatal ages from donors aged 2 to 77; and umbilical cord blood samples. Each sample consists of hematopoietic stem and progenitor cells spanning a spectrum of differentiation states, from uncommitted to committed within 6 distinct lineages. Using this data set, Li et al. [7] identified several age-specific changes in hematopoietic ontogeny. In particular, the independent application of StatOT at each age was used to measure the relative strength with which genes marked commitment to each lineage, which uncovered several age-related transcriptional programs. Here, we demonstrate that gStatOT improves trajectory reconstruction on sparse data by downsampling the samples at each age and comparing the results to age-independent analyses using StatOT and PBA.

Before applying these methods, we applied batch correction to the entire time course to establish smoother ground truth estimates (see SI.5 for details). Due to the significant change in timescale, from weeks to years, between the prenatal and postnatal ages, our model’s regularization coefficients are effectively 0 when bridging these time spans (see the discussion below (14)), and thus they do not encourage mixing them unless *λ* is chosen to be very large. So to avoid unnecessary computation time and allow for *λ* to be chosen appropriately for each scale, we chose to run our tests on the integrated data split into pre- and postnatal time courses, where the former runs from weeks 10 to 23 after conception, and the latter from birth to 77 years. We then established ground truth estimates for cell trajectories and fate probabilities by applying StatOT and PBA to the full samples at each age, following the same procedure for choosing parameters/growth rates used by Li et al. [7] (see SI.5 for details). Each age was then downsampled before applying gStatOT and reapplying StatOT and PBA. gStatOT and StatOT’s estimates on the downsampled data were evaluated against the StatOT ground truth, while PBA was compared to its own ground truth.

On the sparse samples, parameters for each method were fit to a single downsampling of the data, using a weighted average of our marginal, trajectory, and fate probability metrics. See SI.7.3 for all parameter choices used in our evaluations. Each method’s performance was then evaluated on 10 repeated downsamplings of the full time courses.

Figure 6 shows the results of our tests on the downsampled time courses. Panel **A** displays the performance of each method across varying levels of sparsity. We observe that gStatOT is able to maintain an advantage over independent applications of StatOT and PBA in trajectory and fate probability estimation, across multiple levels of sparsity. Notably, it is able to reach the 0.2 TV error threshold with 4 times fewer cells than StatOT and at least 7 times fewer cells than PBA.

**Figure 6.**
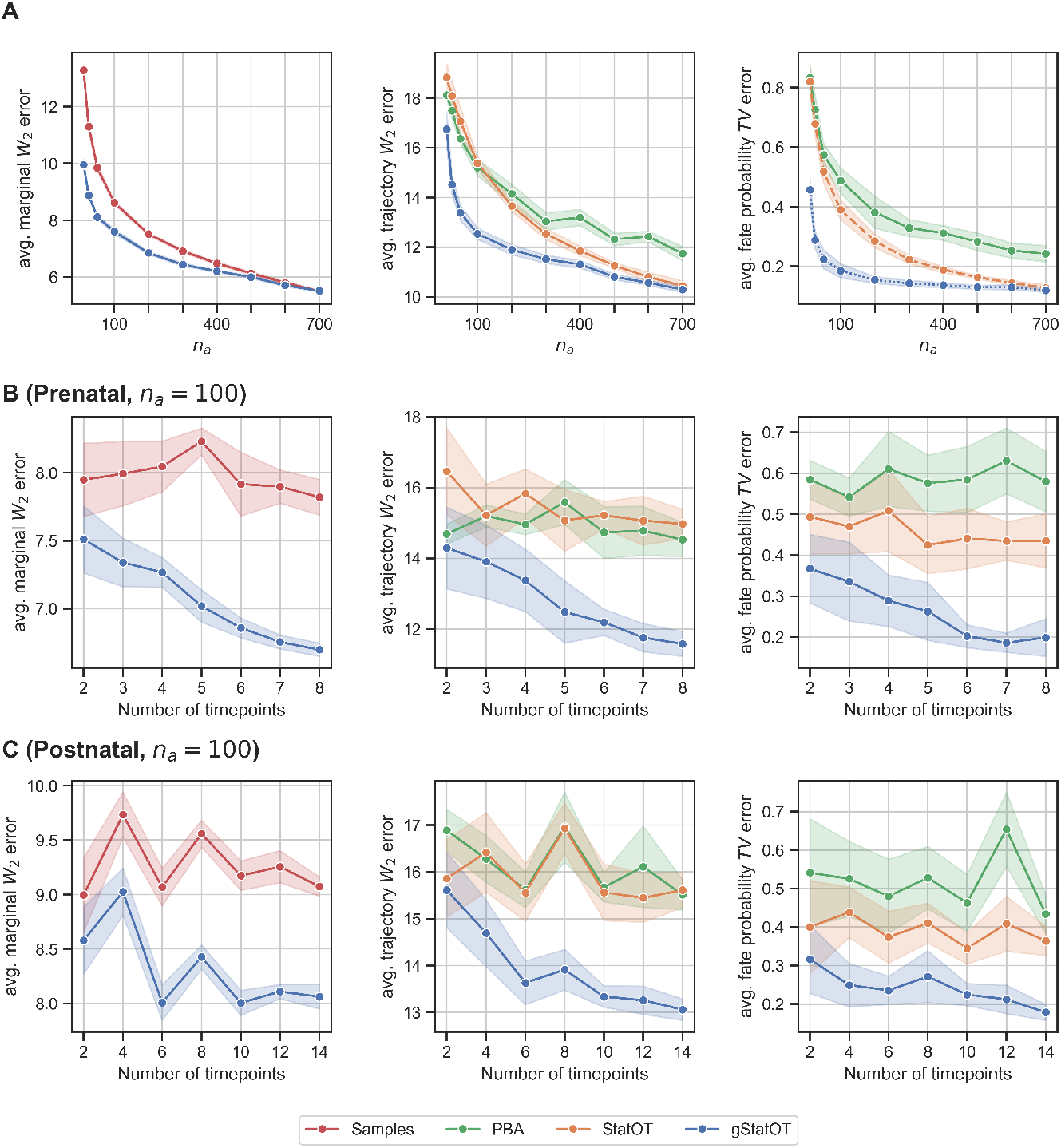
Average performance of each method using time courses with variable sample sizes (**A**) and temporal resolution (**B** and **C**), formed using the hematopoietic time course of Li et al. [7]. **A:** Average performance of each method with varying numbers of cells per age, n_a_ = 10, 25, 50, 100, …, 700. gStatOT took less than 2 minutes to solve on all levels of sparsity using a single NVIDIA H100 80GB HBM3 MIG instance (profile: 3g.40gb). See Figure SI.7 for the exact run times at each level. **B**,**C:** Average performance of each method across all pre- and postnatal ages, when evaluated on time courses with varying temporal resolution, and a fixed number of cells per age (n_a_ = 100). Results in all panels are summarized by the mean and standard deviation of each method’s average performance across age on 10 resamplings of the data.

As with the simulated data, we tested gStatOT’s sensitivity to the number of ages present in the data. To form each time course, we held the initial and final times fixed and added intermediate samples to be as evenly spaced in time as possible. Parameters were tuned for each level of temporal resolution. The average of each metric across the pre- and postnatal time courses is displayed in Panels **B** and **C** of Figure 6. We again observe that the performance of gStatOT improves with an increasing number of time points on both time courses, whereas the StatOT and PBA results remain flat, providing evidence that our method effectively exploits global information across all ages.

To visualize the improved distribution and fate probability estimates obtained with gStatOT, in Figure 7 we display a UMAP embedding of the integrated data coloured by the predicted fate probabilities for each lineage of the week 15 sample, alongside the cell type annotation done by Li et al. [7]. Fate probabilities are saturated based on the strength of the predicted probability, e.g. the uncommitted HSCs are grey, while the cells predicted to be committed to the Lymphoid lineage are a deep orange. Moreover, the points in the gStatOT plot are scaled by the reconstructed marginals, *µ*_*i*_. We can see that gStatOT’s marginals better match the full data distribution, and include the rarer Mono/DC and Baso/Mast lineages that are missing from the StatOT plot. Figure 7 shows the correlation between the cells’ sparsely inferred fate probabilities and StatOT’s ground truth estimates, averaged across all ages. We can see that gStatOT’s estimates are more strongly correlated with the ground truth over all lineages at the *n*_*a*_ = 100 level of sparsity, and maintain this advantage on the less common, Baso/Mast and Megakaryocyte, lineages at the *n*_*a*_ = 300 level.

**Figure 7:**
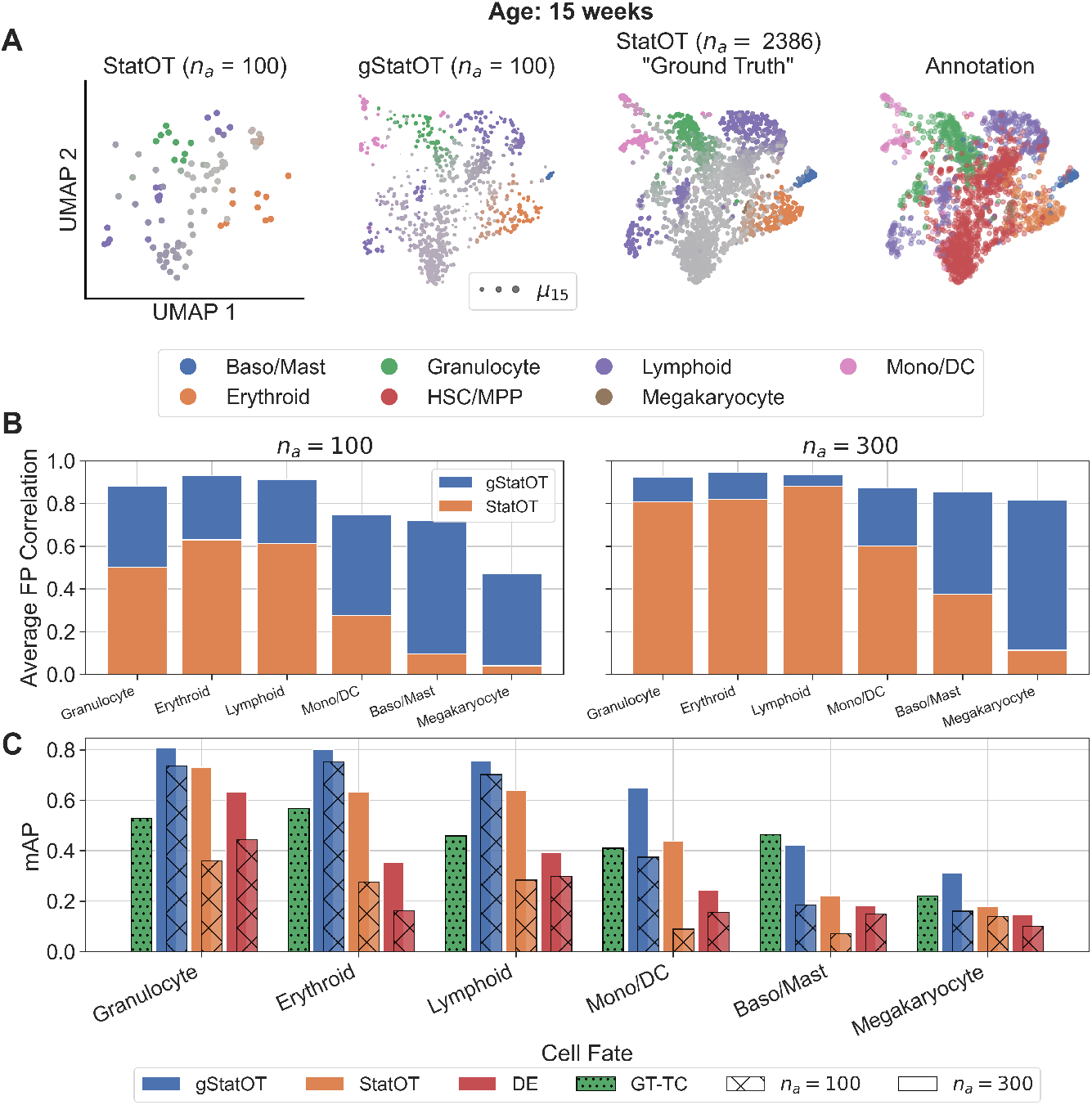
**A:** Visualization of the week 15 sample in the integrated UMAP space, coloured by the predicted fate probabilities for each lineage and method, along with the cell type annotations derived by Li et al. [7]. Fate probabilities are saturated based on the strength of the predicted probability, e.g. the uncommitted HSCs are grey, while the cells predicted to be committed to the Mono/DC lineage are bright pink. The points in the gStatOT plot are scaled by the reconstructed marginals, µ_i_, to show gStatOT’s prediction for the relative number of cells in each state at this age. See Figures SI.5-6 for similar plots for several ages that include each method’s inferred cell flow. **B:** Correlation between the cells’ sparsely inferred fate probabilities and those obtained from the ground truth application of StatOT, averaged across all ages. **C:** A measure of each method’s ability to identify and rank putative driver genes for each cell type across all ages. Each gene was ranked by the correlation between its expression and each lineage fate probability, obtained by gStatOT and StatOT on the sparse samples. The ranked lists at each age were then compared against the top 10 genes identified from our ground-truth StatOT estimates using average precision (AP). The models were evaluated on downsampled time courses with n_a_ = 100 and 300 cells per time point, and the results are summarized by the mean AP (mAP) across all 22 ages. Two baselines are included: GT-TC is the mAP between consecutive ages’ ground-truth gene lists, representing the temporal consistency of these lists, and the DE baseline is the AP of the list of top differentially expressed genes between each cell type and the rest of the data set, at each age. Both baseline results are summarized by the mAP across all ages (see SI.6.4 for details)

A standard analysis downstream of TI is the identification of lineage driver genes. This is accomplished by ranking genes according to their correlation with the inferred lineage fate probabilities [6, 10]. So, we sought to test whether gStatOT’s improved fate probability estimates would translate into better identification of these sets of driver genes from sparse data. To do this, we generated a list of the top 10 putative driver genes for the cell lineages at each age by ranking them according to their expression’s correlation with each lineage fate probability predicted by our previous ground truth application of StatOT. We then evaluated the competency of gStatOT and StatOT in reconstructing these lists at each age from the downsampled time courses using average precision (AP) [39, 40]. Specifically, we computed the AP of each method’s gene list for each fate and age and reported the mean AP (mAP) across all ages in Figure 7 Panel **C** (see SI.6.4 for details).

Our test was evaluated on downsampled data with 100 and 300 cells per time point, using the same model parameters that were fit during the previous evaluations. We observed that gStatOT achieved the best ranking of the top 10 relevant genes across all primary cell types, exhibiting an mAP comparable to or better than StatOT using only a third as many cells. However, both methods struggled on rare cell types such as Baso/Mast and Megakaryocyte. Since gStatOT’s fate probabilities maintain a relatively high correlation with those of the ground truth (Panel **B**) this is likely due to dropouts and sampling noise causing instability in the ground truth lists (measured by the green bar, which compares the ground truth lists at consecutive ages).

## 4 Discussion

We have presented our novel, theoretically motivated method, global StationaryOT, that reconstructs debiased cell trajectories from quasi-static aging single-cell snapshot time courses. We have shown, using both real and simulated data, that gStatOT can correct for data sparsity and provide significantly improved estimates compared to naive schemes that apply TI methods to samples independently. Moreover, the improved, globally supported cell-fate probability estimates enable more accurate identification of the genes involved in the GRN driving cell-fate specification.

Since gStatOT learns the data structure more effectively with increasing sampling rates, we hope it may open the possibility for new experimental designs that favour higher sampling rates over larger individual samples at each age. We speculate that future work on the convergence rate of gStatOT may show that such rapid sampling strategies are optimal when the total number of cells to be processed is fixed.

Our BCA implementation already enables the analysis of sizeable aging time courses, comprising hundreds of cells per time point. However, as we are solving multiple entropic optimal transport (EOT) problems, the method inherits a computational complexity that is quadratic in the number of cells, i.e., *O*(*TN* ^2^). While this scaling has been manageable for only a single sample [5], it presents a significant limitation when applied to time courses with tens of thousands of cells per sample. To address this, low-rank formulations of EOT have recently been used to scale OT-based TI methods to millions of cells [41], and a similar approach could be well suited to extend our method. Alternatively, warm-starting approaches could be explored to reduce the number of required iterations. One such approach could solve the problem on a reduced set of clustered cell states, after which the solution could be used to initialize the problem on the full data set.

Lastly, we note that in this work we have extended previous TI models and methods designed to infer cell trajectories that exclusively traverse gene expression space, and have modelled the effects of aging on these trajectories through the age-dependent epigenetic landscape Ψ. Being restricted to gene expression data prevents us from mechanistically modelling and inferring how epigenetic alterations in each cell drive the observed changes in gene expression and cell fate over age. Future work could extend our method to incorporate epigenetic data, such as chromatin accessibility with scATAC-seq [42], which could not only improve the accuracy of our inferred trajectories, but also allow for the distributional changes in gene expression to be explained by the observed cell states and inferred cell dynamics at previous ages.

## 5 Data and code availability

An implementation of global StationaryOT is available at https://github.com/ColeBoyle/global-stationaryOT. All code for reproducing our results is archived on Zenodo at https://doi.org/10.5281/zenodo.20723235. The hematopoiesis data published by Li et al. [7] is publicly available at https://www.ncbi.nlm.nih.gov/geo/query/acc.cgi?acc=GSE189161.

## 7 Author contributions statement

C.B., E.V., and G.S. conceived of the method. C.B. devised, implemented, and tested the method.

C.B. wrote the manuscript. C.B., E.V., and G.S. reviewed and edited the manuscript.

## 8 Acknowledgments

G.S. is funded in part by a Discovery Grant from the Natural Sciences and Engineering Research Council of Canada, a Canadian Institutes of Health Research Grant, and the United Therapeutics collaboration.

## Supplementary information

**Figure SI.1:**
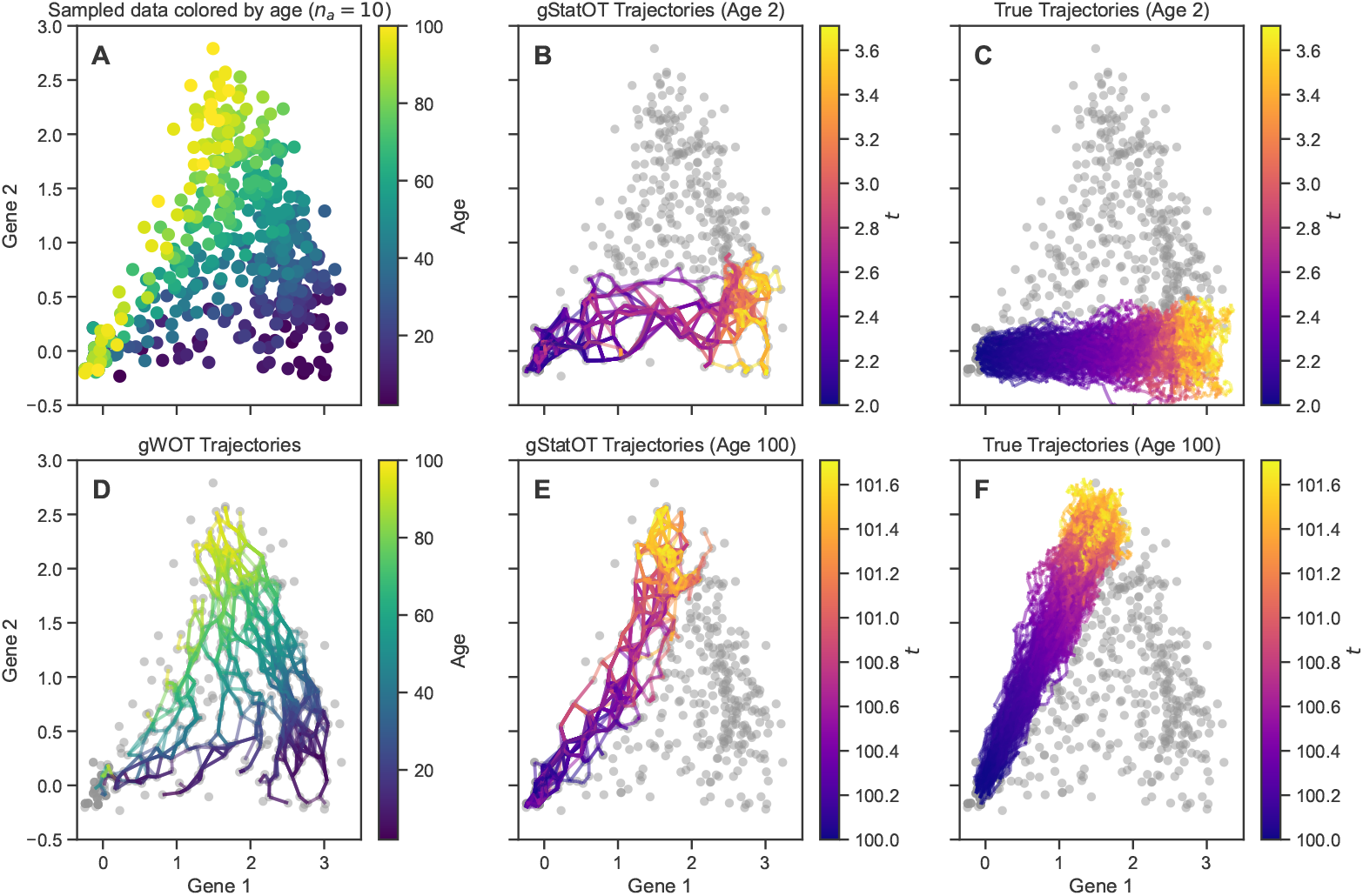
Comparison between trajectory inference with global WaddingtonOT (gWOT)[S1] and global StationaryOT (gStatOT) using time course data. **A:** Data sampled from a simple single lineage, coloured by the age at which it was sampled. Cells start off in growing progenitor states near the origin, and then develop by expressing gene 1. As the system ages, cells begin to express increasing amounts of gene 2 during development. **B:** Cell trajectories inferred by gStatOT at age 2, using all the sampled data. These match the true cell trajectories depicted in Panel **C**, both in that they primarily involve only gene 1 and that they terminate in around 1.6 chronological time units (*t*). **D:** Since gWOT assumes the samples in the time course are the temporal marginals of cell trajectories, the cell paths that it infers traverse the entire aging spectrum. These paths start in the initial, age 2, sample and end in the final, age 100, sample. This makes them perpendicular to the true trajectories, and moreover fails to capture the aging effect in the system. **E**,**F:** In contrast to gWOT, gStatOT leverages the cell growth rates to infer the correct directionality of the paths, while simultaneously utilizing the time course data to infer trajectories that better emulate the ground truth cell paths at every sampled age. The code for generating this simulation and figure can be found at https://github.com/ColeBoyle/global-StationaryOT/extra/gWOT_comparison.

## SI.1 Solving Global StationaryOT

In this section, we show that the discretized version of gStatOT has a unique solution, provided that our cell growth rates satisfy the natural condition that sources and sinks are specified at every age. We then show how we derive and solve the corresponding dual problem.

### SI.1.1 Discretization of gStatOT

Recall that we approximate the full cell state space by the union of our samples, 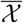. Letting 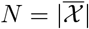 be the total number of sampled cells, with this approximation, 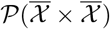 becomes the set of nonnegative *N* × *N* matrices with unit sum whose rows and columns are indexed by the elements of 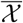. Using this discrete approximation of the state space, our recovery program may be written as

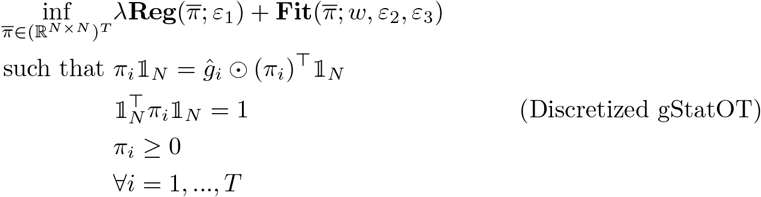

where 1_*N*_ is the vector of 1’s of length *N*. To make the problem easier to solve, we approximate the OT problems in **Fit** and **Reg** by their entropic version, with regularization strength *ε*_1_ = *ε*_3_ = 5 · 10^−3^. See Section SI.2 for more details on selecting the parameters *λ, w*, Δ*t*, and *ε*_2_.

### SI.1.2 Feasibility Condition and the Existence of a Unique solution

Our constraint, which forces a balancing of growth at each age, introduces the feasibility condition that we must have both sources and sinks at each age, or else have neither. gStatOT can naturally satisfy this condition by sharing growth rate information across the time course. Here, we show that this condition is not only necessary but also sufficient for the existence of a unique solution to the discretized gStatOT problem.

Our constraint set is compact and convex (the intersection of a linear and convex constraint, with the unit sum constraint ensuring boundedness), and our objective function is strictly convex and continuous. In order to show the existence and uniqueness of solutions to gStatOT, it suffices to show that this constraint set is non-empty. This occurs exactly when our global sample specifies both source and sink cells at each age. That is, for each *i* = 1, …, *T*, 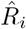 must satisfy the following

There exists *j* such that 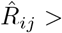 if and only if there exists *k* such that 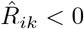.

We note that this also applies to the situation where we have neither sources nor sinks at a given age; however, this case is less interesting since it predicts no cell movement. Moreover, it is unlikely to occur in applications.

*Proof*. Fix some *i* and let 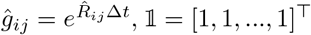. In the discrete setting, our constraints can be written as

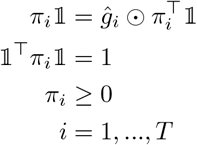

Let 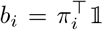. Then there exists a *π*_*i*_ satisfying the constraints if and only if there exists *b*_*i*_ such that ∑_*j*_ *b*_*ij*_*ĝ*_*ij*_ = ∑_*j*_ *b*_*ij*_ = 1, *b*_*ij*_ *≥* 0. That is, there must exist a positive solution to the following linear system

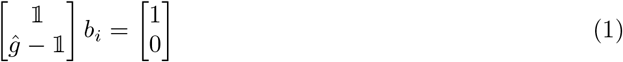

Farkas’ lemma [S2] implies that *b*_*i*_ exists if and only if there does not exist *y*_1_, *y*_2_ ∈ ℝ such that

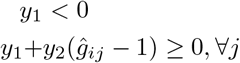

Clearly, if *ĝ*_*i*_ *>* 1 we may choose *y*_2_ = 1 and − min_*j*_(*ĝ*_*ij*_ − 1) *< y*_1_ *<* 0, and a similar situation exists if *ĝ*_*i*_ *<* 1. Whereas, if there exists *j, l* such that *ĝ*_*ij*_ *<* 1 and *ĝ*_*il*_ *>* 1 then we would need

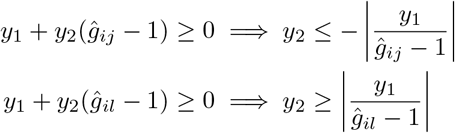

These conditions would imply that *y*_2_ = 0, but *y*_2_ can’t be 0 since this would contradict the first condition that *y*_1_ *<* 0. Finally, if *ĝ*_*i*_ ≡ 1, the conditions above imply that *y*_1_ *<* 0 and *y*_1_ ≥ 0, thus no solutions exist. Therefore, Farkas’ lemma implies that a non-negative *b*_*i*_ solving Equation 1 exists if and only if our stated condition holds.

### SI.1.3 Derivation and solving of the dual problem

Converting each of the OT problems in **Reg** and **Fit** to their dual forms and converting our growth constraint into a Lagrange multiplier, we may write the discretized gStatOT problem as

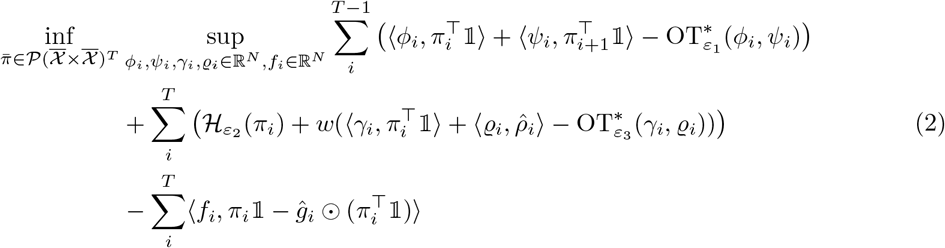

where

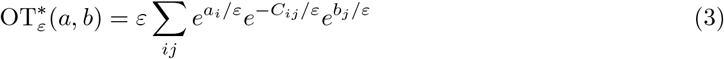

is the Legendre transform of OT_*ε*_. Let 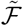 denote the objective function in (2). Since 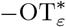 is jointly concave in its arguments, 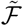 is concave in *ϕ, ψ, γ, ϱ* for fixed 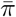. Moreover, 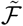 is convex in 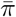 when all other arguments are fixed since ℋ_*H*_ is convex and the maps *π* ↦ ⟨*a, π*1⟩, *π* ↦ ⟨*a, π*^⊤^ 1⟩are linear for all *a* ∈ ℝ^*N*^. Thus, since 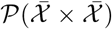 is convex and compact, we may apply Sion’s minimax theorem to swap the inf and sup

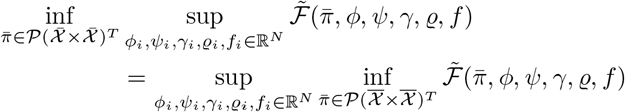

The last step is a standard application of the Fenchel-Rockafellar (FR) duality theorem to remove our final constraint. Let *A* be the linear map that sends 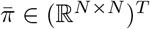 to the sums of its elements, 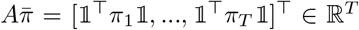. Letting 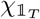 be the indicator for the vector of ones of length *T*, and noting that 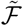 is only finite when 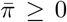 (and we know such a solution exists because of the previous result), we may write

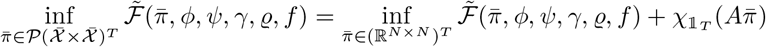

Next, holding all dual variables fixed, the Legendre transform of 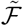 and 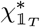 is given by

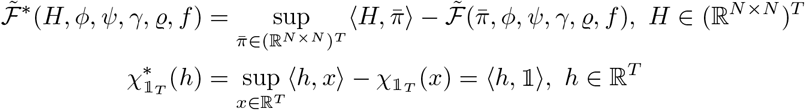

Now we can apply FR [S3, Theorem 4.4.3],

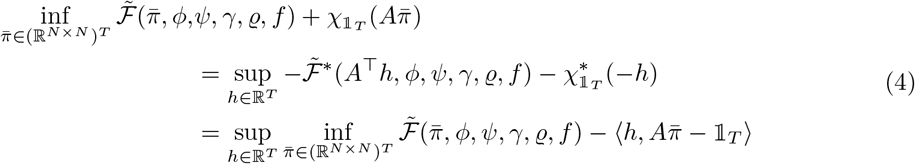

The inner minimization problem is solved by simply setting the gradient of 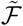 in 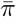 to zero, holding all else fixed. By doing so, we obtain the optimal 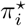

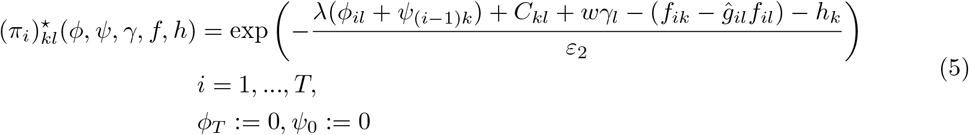

Substituting 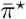 into (4), we obtain the dual problem of discrete gStatOT

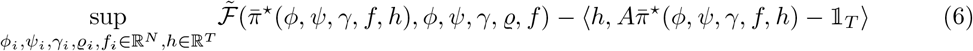

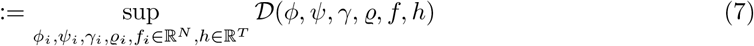

### SI.1.4 Implementation

#### SI.1.4.1 Block coordinate ascent solver

The total gradient of *D* can be manually computed, and off-the-shelf gradient based solvers can be used to solve (7). However, this requires evaluating the exponential in (5), which can lead to numerical instability if *ε*_2_ is small. Instead, analogous to the algorithm used to stabilize traditional entropic OT problems [S4], we use a block coordinate ascent (BCA) approach [S5] that allows us to exploit the log-sum-exp trick to stabilize exponential evaluations.

In particular, we update each block of dual variables, *ϕ, ψ, γ, ϱ, f, h*, sequentially by maximizing *D* with respect to the block being updated, while holding all other blocks fixed. The updates for *ϕ, ψ, γ, ϱ* and *h* are given by closed-form expressions, as ∇_*z*_*D* = 0 can be solved analytically for *z* ∈ {*ϕ, ψ, γ, ϱ, h*}^∗^. All of these expressions can exploit the log-sum-exp trick to avoid evaluating exponentials. However, the update for *f* is not given by a closed-form expression, since 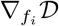 is equal to our growth constraint and 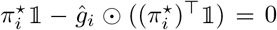 is a highly coupled set of nonlinear equations in *f*_*i*_. Therefore, we update *f* by performing a single step of the LBFGS algorithm [S6] on 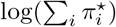 with respect to *f*. This step is equivalent to maximizing *D* with respect to *f*, but allows us to again exploit the log-sum-exp trick to stabilize the computation.

We repeat this process until convergence, which we determine by evaluating the relative duality gap. If *D* is the dual objective and *P* is the primal objective, we compute the duality gap as *G*(*P, D*) = *P* − *D*, and the relative duality gap as 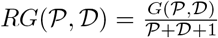. We stop once *RG*(*P, D*) *<* 10^−3^, which we find to be sufficient for convergence in practice. Computing the objectives does require evaluating the exponential in (5), so we only check for convergence every 100 iterations and proceed even if the duality gap is not computable, as early instability in these evaluations doesn’t affect the rest of the algorithm.

We implemented this algorithm in Python using the JAX library [S7] and used the optax [S8] implementation of LBFGS for updating *f*, allowing it to be efficiently executed on GPUs. The code for this implementation can be found at https://github.com/ColeBoyle/global-StationaryOT.

## SI.2 Parameter selection

gStatOT has four main parameters: *λ*, which controls the between sample regularization strength; *w*, which controls the strength of the fit to the sampled data at each age; *ε*_2_, which controls the strength of the entropic regularization in **Fit**; and Δ*t*, which controls the time step of our splitting approximation and represents the timescale at which we expect cells to change state.

*w*: We find that on sparse data gStatOT is robust to the choice of *w*, and so we simply set it to 1 for most of our analyses. As the sample size increases, to match the dynamics inferred by StatOT we must increase *w*. For example, with the *n*_*a*_ = 500, 600, and 700 time courses in Figure 6, we set *w* to 10.

*λ*: The choice of *λ* depends on the sampling rate and sample size, and so must be tuned based on the sparsity of the specific dataset. *λ* should decrease with increasing sample size and sampling rate. However, as noted in the text, *λ* contains an implicit estimate of the aging rate, *λ* ∼ 1*/c*, and so it may be desirable to choose *λ* larger if the aging rate is expected to be slow, regardless of the sampling rate. In practice, we find that an initial value of *λ* between 5 and 20 works well for our tested datasets, where our quantitative results were relatively stable across this range.

*ε*_2_: Based on the connection between diffusion processes and optimal transport, *ε*_2_ is theoretically equal to *σ*^2^Δ*t*. In practice, we simply take *ε*_2_ to represent the expected noise in the data and tune or choose Δ*t* separately. A common heuristic for finding this noise level is to adjust it so that the total predicted fate probability for each lineage matches the proportion of cells previously identified to be a part of that lineage (e.g., with clustering and marker genes) [S9, S10], see Figure SI.4 for an example of this approach for choosing *ε* using StatOT and PBA’s diffusion parameter *D*.

Δ*t*: Since our splitting approximation improves with decreasing Δ*t*, it is ideally chosen to be as small as possible. However, this can lead to overly flat cell growth, *ĝ*_*i*_, which disincentivizes cell movement. So we recommend taking it to be large enough so that cell trajectories converge to sink states within a reasonable number of iterations. The value 0.25 worked well as a default value for StatOT, and we used this value for our hematopoiesis study.

## SI.3 Fate probability estimation

### SI.3.1 Computing fate probabilities with flux rates

To compute the fate probabilities of cells, we expand upon the method used in the original StatOT paper [S10], by incorporating the flux rate of cells dying or leaving the system instead of simply treating every sink cell as an absorbing state.

We use a similar approach to that of Weinreb et al. [S9]. In particular, for each of the *K* lineages or cell types, *l*_*k*_, that are present in our system, we build a sink matrix *S* ∈ ℝ^*N*×*K*^, where *N* is the number of cell states and for cells labeled as being part of lineage *l*_*k*_, we take *S*_*jk*_ to be the probability that the cell 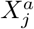 leaves the system as a type *l*_*k*_ within the period of time Δ*t*, namely, 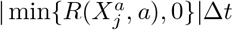. After row-normalizing our couplings *π*_*i*_ we augment them with *S* to form a new transition matrix, *P*_*i*_, that describes the probability of transitioning from one cell state to another, or leaving the system in a particular lineage over the period of time Δ*t*. In particular, we have

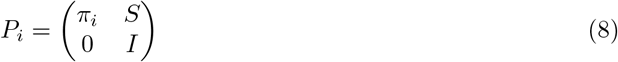

After another row normalization, we interpret *P*_*i*_ as the transition matrix of a Markov chain, and proceed to compute the fate probabilities of cells by solving for the absorption probabilities of this Markov chain as previously done (see supplementary section 1.2 of [S10]).

### SI.3.2 Stabilizing fate probability estimation

As with other methods that estimate cell dynamics with Markov chains [S9, S11, S10] our ability to estimate the cell dynamics does not always imply that we are able to stably compute cell fate probabilities. For the OT based methods, this occurs when one wishes to use small *ε* values which can lead to sparse transition matrices, and thus disconnected cell states that are unable to reach any of the sink states. In practice, this can occur if the data contains outlier cell states that are predicted to be rare and reachable only by fully random motion and not the natural flow of cells. In this case, the resulting transition matrices may be ill-conditioned for solving for cell fate probabilities.

Other methods [S11, S10] have addressed this limitation by increasing the base noise level by perturbing the transition probabilities a small amount after they have already been fit to the data. Here, we employ an alternative method that seeks to remove these outlier states before computing the fate probabilities for the rest of the cells. In particular, if *X*_*j*_ is an outlier state, then little to no mass should flow through this state, hence, our inferred coupling *π* will assign very low probabilities to points (*X*_*j*_, *X*_*k*_) or (*X*_*k*_, *X*_*j*_) for *k* ≠ *j*. That is, setting mass in_*j*_ = ∑_*k*_ *π*_*kj*_ − diag(*π*)_*j*_ and mass out_*j*_ = ∑_*k*_ *π*_*jk*_ − diag(*π*)_*j*_, we will have that mass flow_*j*_ = mass out_*j*_ + mass in_*j*_ is very small. Thus, we can detect and remove outlier states by simply pruning states with low mass flow. To implement this, we compute a mass flow distribution by computing mass flow_*j*_ for each cell state^†^, and then remove those states with mass flow below the *q*-th percentile of this distribution. *q* is initially chosen to be 0, but if we fail to compute fate probabilities due to ill-conditioning or poor convergence, we increase *q* by 0.01 and repeat this process until we are able to compute fate probabilities, or until *q* reaches a maximum value of 0.1. In practice, we find that this method allows us to stably compute fate probabilities for a wider range of *ε* values, and thus obtain more accurate estimates of cell dynamics when the overall noise level in the data is low, but our samples still contain outliers. We employ this strategy when computing fate probabilities with both gStatOT and StatOT. Lastly, we note that this only works when using the coupling *π*. Once *π* is row-normalized to form the transition matrix we lose the ability to compare the mass flow of different states.

## SI.4 Simulation Details

### SI.4.1 General model application details

#### gStatOT and StatOT

For both our simulations, gStatOT and StatOT’s Δ*t* parameter were set to the simulation time step *dt*. Fate probabilities were computed as described in Section SI.3. The StatOT algorithm was implemented using the Optimal Transport Tools (OTT) Python library [S12]. In order to provide the best possible competitor to gStatOT, we used the log-domain stabilized version of the Sinkhorn algorithm implemented in OTT, which provides greater stability in low *ε* settings. For greater stability, both methods were given cost matrices scaled by the mean of their entries. Moreover, both methods were given the true growth rates of cells at the age they were sampled. For an analysis of the sensitivity of both methods to growth rate estimation error, see Figure SI.2. Other parameters were selected via the parameter sweeps described below.

#### PBA

To apply PBA [S9], we ran the `PBA_pipeline.py` script provided by the authors at https://github.com/AllonKleinℒab/PBA. We made slight modifications to the script to allow us to run it within our testing framework, mainly to make sure it didn’t crash when it failed to compute fate probabilities at a particular age. In such cases, we reported the fate probability metric as 1, which is the maximum value it can take. We also modified the script to allow us to save the transition matrices that PBA constructs so that we could sample individual cell trajectories to use for our trajectory metric. As with gStatOT and StatOT, we gave PBA the true growth rates of cells at the age they were sampled. We set the lineage specific flux rates, *S*^*a*^ ∈ ℝ^*N*×*K*^, *that PBA u*ses to compute fate probabilities, to be the death rate of sink cells in that particular lineage, i.e., 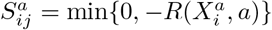 if *X*_*i*_ is a part of lineage *l*_*j*_. The value of *k* used to construct the *k*-NN graph, and diffusion level *D*, that PBA uses to construct its transition matrices were selected via the parameter sweeps described below. The exact code we used is archived at https://doi.org/10.5281/zenodo.20723235.

### SI.4.2 Synthetic data: Aging epigenetic landscape

For our first simulation, we generated a system that evolves according to Equation 3 in the main text. We used the potential

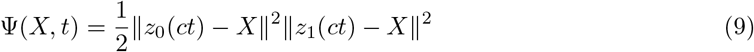

where *c* = 5 × 10^−3^*/*day is our aging rate, and *z*_0_, *z*_1_ are well locations

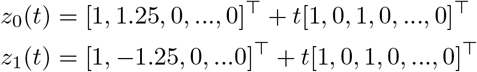

representing the location of the terminal cell states at the age *a* = *ct*. The following cell division rate, *α*, and death rate, *β*, were used

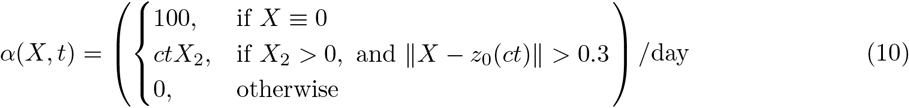

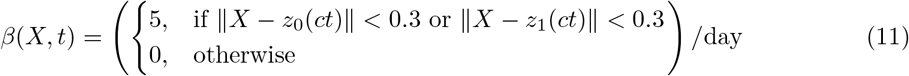

Lastly, the diffusion coefficient was taken to be *σ*^2^ = 0.1.

To generate an aging time course from this system, we started with a population of 5 fixed cells (which do not feel the effects of drift or diffusion) located at the origin, and simulated this system for 99 days using the Euler-Maruyama method with a time step of *dt* = 0.01 days. Division and death were applied at each iteration by adding (if *R*(*x, a*) = *α*(*x, a*) − *β*(*x, a*) *>* 0) or removing (if *R*(*x, a*) *<* 0) a cell at position *x* and age *t*, with probability |*R*(*x, a*)| *dt*. We generated an aging time course by sampling all cells from the simulation every day starting at day 2. This time course was then subset using the various sampling rates listed in Figure 4.

For each sample in the time course, a ground truth distribution of cell trajectories was generated by simulating 500 individual cell trajectories, each starting from a cell sampled with weight proportional to its division rate, max {*R*(*x, a*), 0}. Each trajectory was run until the cell died, revealing an empirical estimate that the average cell lifespan was approximately 3.5 days at day 2 and decreased to 2 days at day 99, due to the increase in shorter cell trajectories resulting from the growth increase in the +2 lineage. So we truncated all trajectories to 2 days for comparison with each method’s reconstructed trajectories.

Estimates of the ground truth fate probabilities were obtained by simulating 500 trajectories starting from each sampled cell and computing the fraction of trajectories that ended in the +2 and −2 lineages, respectively.

Time courses with each of the specified sampling rates in Figure 4 were downsampled to *n*_*a*_ = 10 cells per age before applying gStatOT to the full time course and the methods StatOT and PBA to each individual time point.

For each sampling rate, we performed a sweep over each method’s parameters and selected those that minimized the mean of the marginal, trajectory, and fate probability metrics (see Section SI.6 for a detailed discussion of these metrics). To evaluate the trajectory metric, 2 day long (i.e. 200 step) cell paths were sampled from each of the method’s transition matrices. Before taking the mean over all the metrics, we scaled each metric by its minimum and maximum values over all method and parameter choices. As discussed in Section SI.3, certain parameter values may lead any of the methods to produce transition matrices that are ill-conditioned for computing fate-probabilities; moreover, the lack of sink states at some ages makes it impossible for StatOT and PBA to compute these probabilities. If a method failed to compute fate probabilities at a certain age we set the total variation metric to its maximum value of 1. Trajectories that were shorter than 2 days were padded with their final state to reach the length required for comparison.

The mean metric and optimal parameters for each method and sampling rate are shown in Section SI.7.1. The parameters used for the results depicted in Figures 3 and 4 in the main text are highlighted in red. If a method failed to produce fate probabilities at any age for that parameter choice we highlighted the mean metric value in orange.

To produce the results in Figure 4, the methods were run using the optimal parameters for each sampling rate, on 10 reseedings of the simulation and time course generation process. The results were summarized by taking the mean and standard deviation of each metric across the 10 runs.

Code for generating the simulation data and figures 3 and 4 in the main text can be found at https://github.com/ColeBoyle/global-StationaryOT/.

### SI.4.3 Synthetic data: Loss of cellular identity over age induced by epigenetic erosion

To develop a simulation that more closely resembles a real aging developmental system, we employed the gene regulatory network (GRN) simulator BoolODE [S13], which simulates realistic scRNA-seq data resulting from the dynamics of a user-specified Boolean GRN with specified gene interaction strengths. However, BoolODE does not support simulating either cell growth or time dependent interaction strengths, and so we used BoolODE to translate the GRN in Figure 5 Panel A into a stochastic ODE model and modified the simulation to support both of these components.

In more detail, BoolODE takes a Boolean GRN and a list of gene interaction strengths, *S*, as input and outputs a model for cell velocities in the gene and protein expression spaces. Specifically, letting a cell state be denoted by *X*_*t*_ = (*G*_*t*_, *P*_*t*_), where *G*_*t*_ denotes the list of genes and *P*_*t*_ the list of associated proteins, BoolODE outputs a velocity vector, *V* (*X*_*t*_, *t, S*), that describes the deterministic part of the dynamics of *X*_*t*_. Normally, BoolODE would then simulate cell trajectories evolving according to the SDE

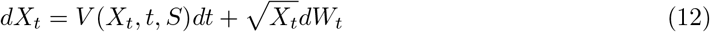

with boundary conditions to ensure that gene and protein expression remain non-negative.

To simulate the loss of cellular identity over age that results from epigenetic erosion, we modified the regulation strength, *S*_*B*1↔*B*2_ ∈ *S*, between the lineage driver genes, *G*_*B*1_ and *G*_*B*2_, so that it decreased linearly with age from 0.9 at age 0 to 0.5 at age 150. We incorporated cell growth by giving cells rates of division and death that depended on the abundance of the proteins *P*_*g*_ and *P*_*d*_, that are coded for by the growth gene *G*_*g*_ and apoptosis gene *G*_*d*_ respectively. Specifically, cells with more than 15 units of *P*_*g*_ were given a division rate of 2 per (chronological) age, while cells with more than 15 units of *P*_*d*_ were given a death rate of 1 per age.

With our modified *S*, which now depends on age, we simulated (12) for 150 ages with the Euler-Maruyama method with a time step of *dt* = 0.01 ages, starting with 5 fixed cells initialized with 1 unit of *G*_1_ and *G*_*g*_. Cell growth was simulated using the same method as in the previous section. We generated a time course with 28 time points by sampling all cells from the simulation every 5 ages starting at age 15 (the chronological age at which the system had reached a near equilibrium state). We then induced dropouts in the data following BoolODE’s procedure. Namely, genes with expression below the median expression level of all cells at a particular age were set to 0 with probability 0.5.

The data was then labeled by assigning cell types for each of the branches, *B*1, *B*2, and *B*1 + *B*2, based on whether a cell exclusively expressed one of the terminal lineage markers *G*_*B*12_ or *G*_*B*22_ above a threshold of 1, or else expressed both above this same threshold. Lastly, cell fate probabilities were estimated by simulating 500 trajectories starting from each cell and computing the fraction of trajectories that ended in each of the three branches.

This simulation can be generated by downloading the BoolODE code from https://github.com/murali-group/BoolODE and running the script at https://github.com/ColeBoyle/global-StationaryOT/extra/data_preprocessing/sim/GRN_simulation. The exact BoolODE code we used is archived at https://doi.org/10.5281/zenodo.20723235.

To produce the results in Panels **D** and **E** of Figure 5, the time course was downsampled to *n*_*a*_ = 25 cells per age before applying gStatOT, StatOT, and PBA. All methods were given the true growth rates of the sampled cells, *R*(*X, a*), and gStatOT and StatOT were given the integration time step *dt* as their Δ*t* parameter. All other parameters were tuned to optimize for fate probability estimation, with the results and optimal choices shown in Section SI.7.2. With these parameters, the methods were run on new downsampled data from 10 reseeded simulations, and the results were summarized by taking the mean and standard deviation of each metric across the 10 runs.

**Figure SI.2:**
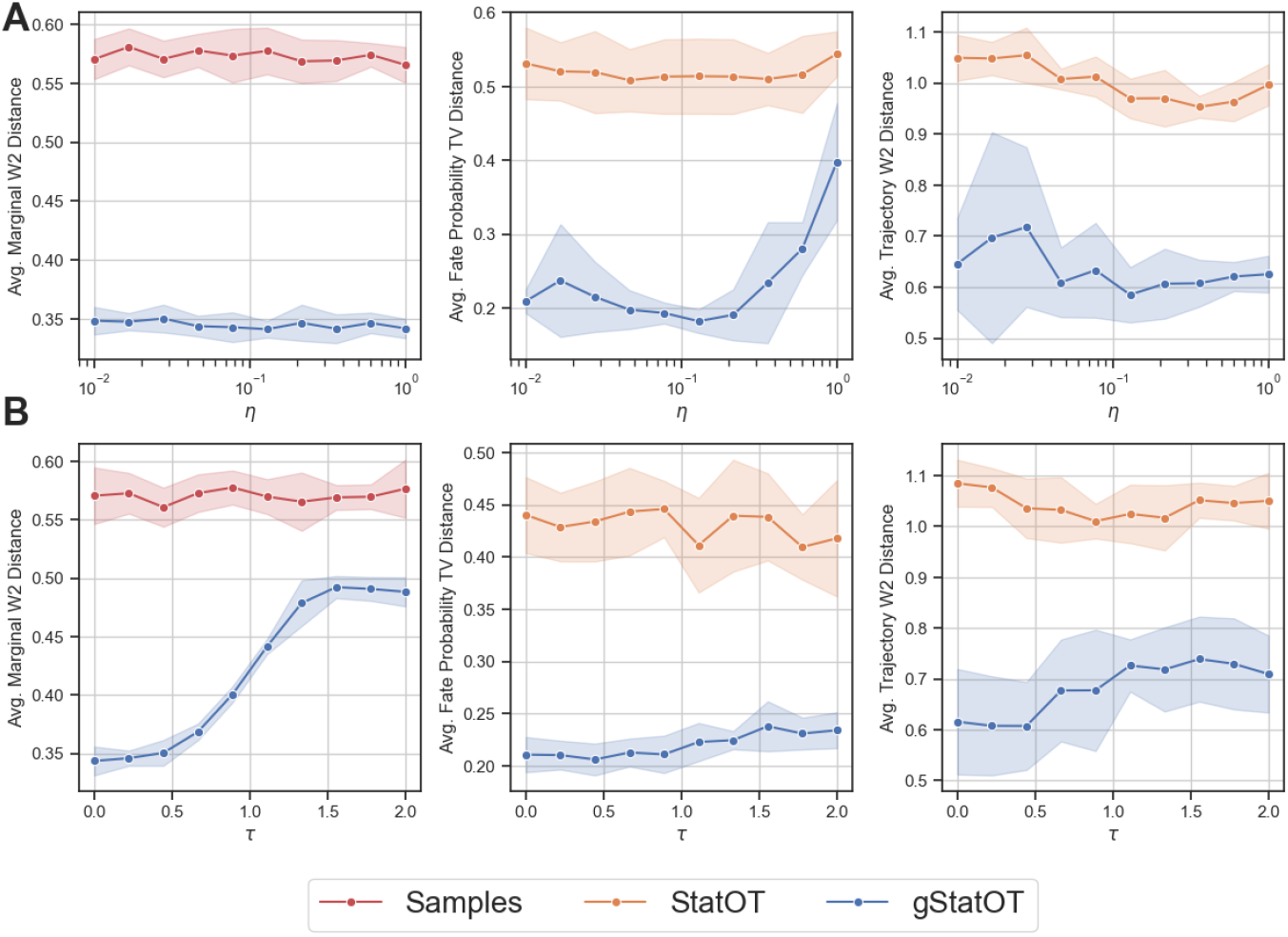
gStatOT’s sensitivity to growth rate noise and misspecification. We tested gStatOT’s outputs on the simulated time course (with *T* = 25 samples of *n*_*a*_ = 10 cells) presented in Figures 3 and 4 of the main text. We used the same parameters as in the main text, but with noisy (Panel **A**) and misspecified (Panel **B**) growth rates. In Panel **A**, we gave gStatOT and StatOT growth rates subject to both additive and multiplicative noise at various levels. We used multiplicative noise to account for the large disparity in growth rates between cell states, and additive noise for the majority of cells that have a true growth rate of 0. Specifically, we set each cell’s growth rate to 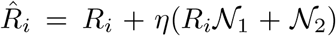, where *R*_*i*_ is the true growth rate, *N*_1_ and *N*_2_ are independent standard normal random variables, and *η* is a noise level parameter selected to be 10 evenly spaced values in the logarithmic space between 0.01 and 1. We found gStatOT’s performance to be relatively robust to this noise, with the fate probability error significantly deviating only once the noise level was on the same order of magnitude as the spatial noise 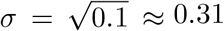. In Panel **B**, we again kept all parameters the same but gave both methods misspecified growth rates by giving an increasing number of uncommitted cells the 100 per day division rate of the fixed progenitor cells in our simulation. In particular, we defined these uncommitted cells to be ones with absolute Gene 2 expression less than 0.25, and Gene 1 expression less than *τ*, with *τ* selected to be 10 evenly spaced values between 0 and 1. Here, we found gStatOT’s fate probability estimates to be stable, however, the marginal error steadily increased with *τ*, and a similar but smaller trend was observed in the trajectory error. This change in the marginal error is expected due to the significant imbalance in growth rates between the source and sink states. In the previous test, the noise was applied uniformly and so the relative difference in growth between source and sink states remained similar. In contrast, the large number of high growth cells forces the method to decrease the mass on these states in order to satisfy the equilibrium constraint, leading to erroneous marginals supported only on committed cell states. However, even with this misspecification, gStatOT’s marginals were still seen as improvements over the raw samples. With both of the tests, we did not find significant trends in StatOT’s performance, as its estimates were already quite poor and variable to begin with. Code for generating this figure can be found at https://github.com/ColeBoyle/global-StationaryOT/extra/.

**Figure SI.3:**
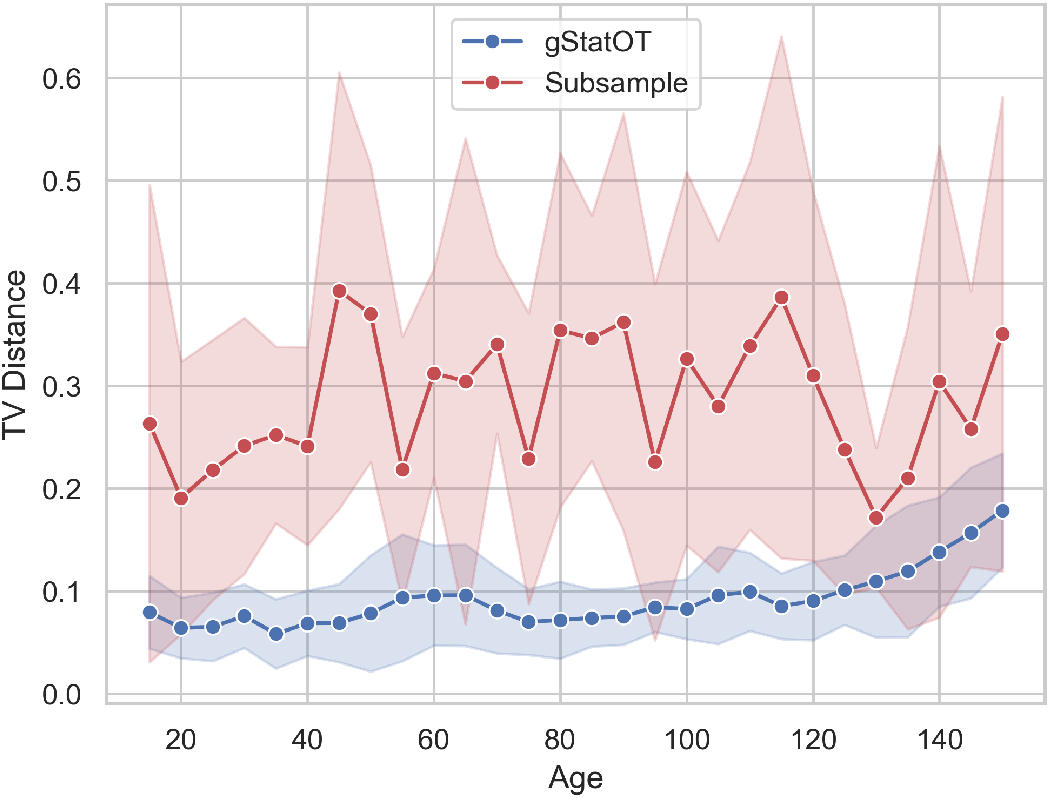
The total variation distance between the proportions of cell death in the simulation and those estimated by gStatOT and the subsampled data (see (13)). The results are summarized by the mean and standard deviation of this distance across 10 reseeded simulations.

To produce the results in Panel **F** of Figure 5, at every age we computed the total number of cells predicted to die in each branch over the time period *dt*, with each of the subsampled data, gStatOT, and the full simulated data. For gStatOT we used the parameters determined above. In particular, for all sink states, 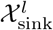, in each lineage, *l* ∈ {B1, B2, B1+B2}, we computed at every age, *a*,

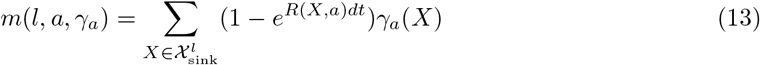

where *γ*_*a*_ was taken to be the subsampled data, 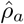, gStatOT’s reconstructed distribution, *µ*_*a*_, and the full simulation samples, *ρ*_*a*_. At each age we plotted the proportion of cell death in each lineage, 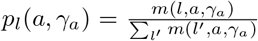, for each of the three *γ*_*a*_’s. To quantify this comparison, we computed the total variation distance from the proportions estimated by the subsampled data and gStatOT to those estimated by the full simulation samples. The results are shown in Figure SI.3.

## SI.5 Hematopoiesis preprocessing and parameter selection

We followed the preprocessing steps outlined in Li et al. [S14] to integrate their hematopoiesis data. Namely, 2000 variable genes were found on each log normalized sample independently, then Seurat’s [S15] (V5.5) `IntegrateData` function was used to integrate the data after finding integration anchors using `FindIntegrationAnchors` with the default parameters. All integration and preprocessing scripts are available at https://github.com/ColeBoyle/global-StationaryOT/extra/data_preprocessing/hematopoiesis. A 30-component PCA embedding was then computed on the integrated data. We then split the data into pre- and postnatal time courses as described in the text. The PCA embeddings of cells in each time course were then passed to both StatOT and PBA to establish ground truth estimates. We followed a similar procedure to that of Li et al. [S14] to fit the methods to the data at each age. In particular, we used the same sink states determined by Li et al. [S14] using marker genes for each of the 6 terminal lineages. For StatOT Δ*t* was taken to be the default parameter 0.25, and the sink states were given a growth rate such that all mass is removed from these states within the period Δ*t*. All other cells were given a small positive growth rate consistent with normalizing *ĝ*_*i*_ after sink growth rates have been accounted for. We chose *ε* = 0.04 so that the total fate probability across all cells, for each of the 6 lineages, was roughly equal to the fraction of cells in the data that were labeled as being part of that lineage by Li et al. [S14] (see Figure SI.4). For PBA, we took *k* = 20 for its *k*-NN parameter and gave sink states flux rates, *R*_*i*_, equal to 1*/D*, where *D* is an estimated diffusion constant. All other cells were given a flux rate such that ∑_*i*_ *R*_*i*_ = 0. *D* = 1 was then selected the same way as *ε* for StatOT. As in [S14], we found that the methods produced fate probability estimates that were highly correlated with each other (see Figure SI.4). Separately for each method and age, we estimated the 80^*th*^ percentile of the hitting time of sink states by sampling cell trajectories from the method’s transition matrix. We then sampled cell trajectories of this length for use in the trajectory metric described in Section SI.6.

These methods were then reapplied to the downsampled data, as described in the text, and parameter sweeps were performed to select optimal parameters for each method, time course, sparsity level, and temporal resolution. gStatOT/StatOT were given the same growth rates and Δ*t* value used when computing the StatOT ground truth on the full samples. PBA was given the same flux rates used when computing its ground truth on the full samples. As with the simulated data, we picked parameters using the average of our evaluation metrics, however since we are using StatOT as a proxy for the ground truth, when choosing gStatOT’s parameters we weighted the marginal metrics twice as high as the others. SI.7.3.1 and SI.7.3.2 show the results of our parameter sweeps for variable sample sparsity and temporal resolution. The parameter values used in Figures 6 and 7 are highlighted in red. Note that for the *n*_*a*_ = 500, 600 and 700 samples sizes we took gStatOT’s *w* parameter to be 10, whereas in all other experiments it was set to 1. See Figure SI.7 for the time it took to solve gStatOT for each sparsity level in Figure 6.

In Figure 7 **A**, fate probabilities were plotted on a UMAP of the entire integrated data set, with fading saturation to indicate the magnitude of each estimate. The points in gStatOT’s plots were scaled by its reconstructed marginals, *µ*_*i*_. Figure SI.5 and Figure SI.6 show similar plots with the inferred flow fields of gStatOT and StatOT at several prenatal and postnatal ages, respectively. These flow fields were computed by taking the row-normalized couplings, 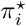, and computing the average velocity of each cell state as 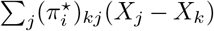, where *X*_*k*_ is the UMAP embedding of the *k*^*th*^ cell state. The flow fields were then plotted as streamlines using Matplotlib’s [S16] `streamplot` function.

For Figure 7 **B**, we computed the Pearson correlation between the sparsely estimated fate probabilities of gStatOT/StatOT and those computed by StatOT for the same cells using the full data. The results were averaged across all ages and reported for each cell lineage.

See Section SI.6.4 for the details of how we computed and compared the lineage driver gene list of each method to produce Figure 7 **C**.

**Figure SI.4:**
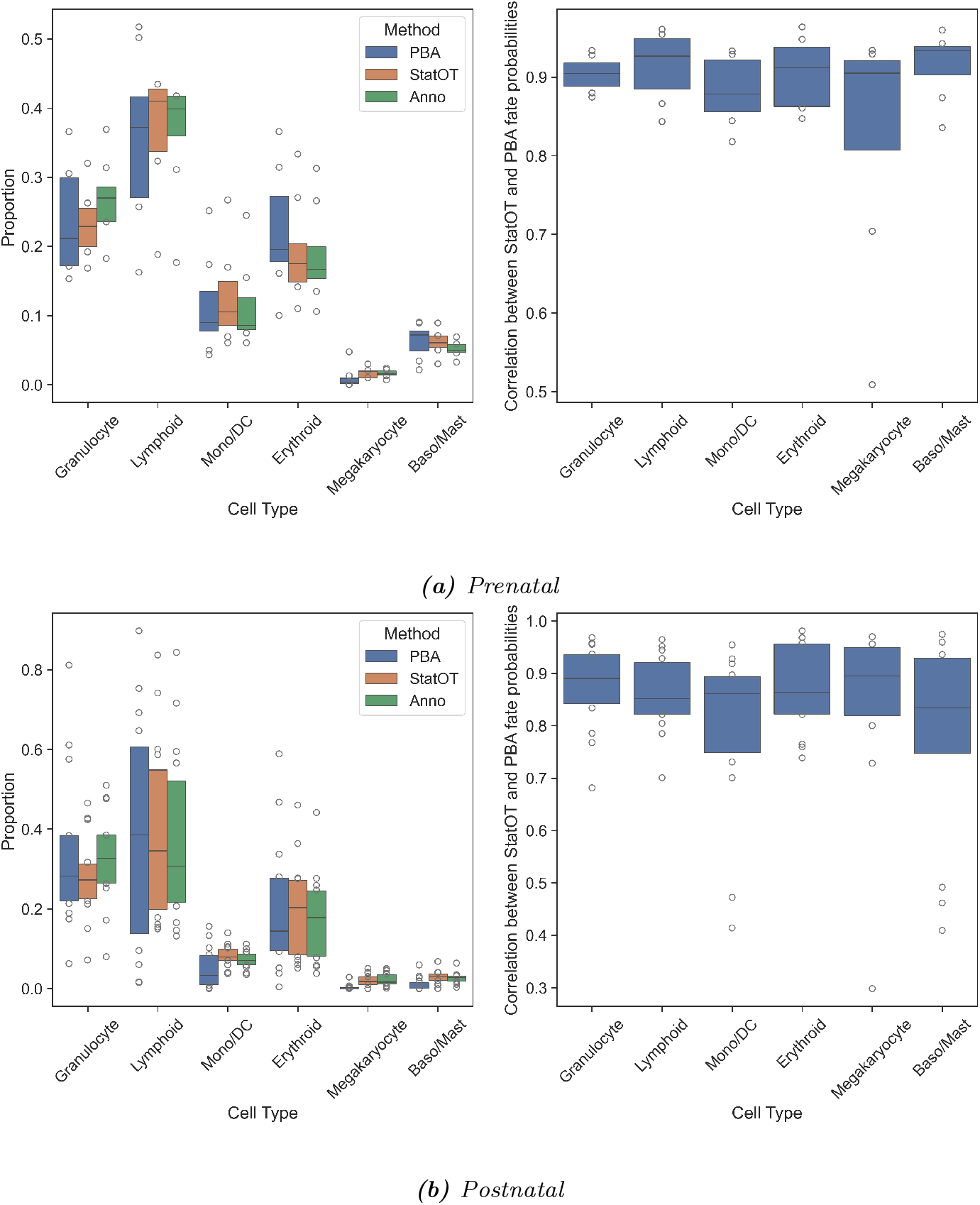
**Left:** Fate probability proportions, for each lineage estimated by StatOT and PBA on the full data, and the proportion of cells in the data annotated as being part of each lineage. Each proportion is the sum of all the cells’ probability of ending in that lineage. StatOT’s ε and PBA’s D parameters were chosen so that each fate probability proportion roughly matched the annotated proportions. **Right:** The Pearson correlation between the fate probability estimates of StatOT and PBA on the full data.

**Figure SI.5:**
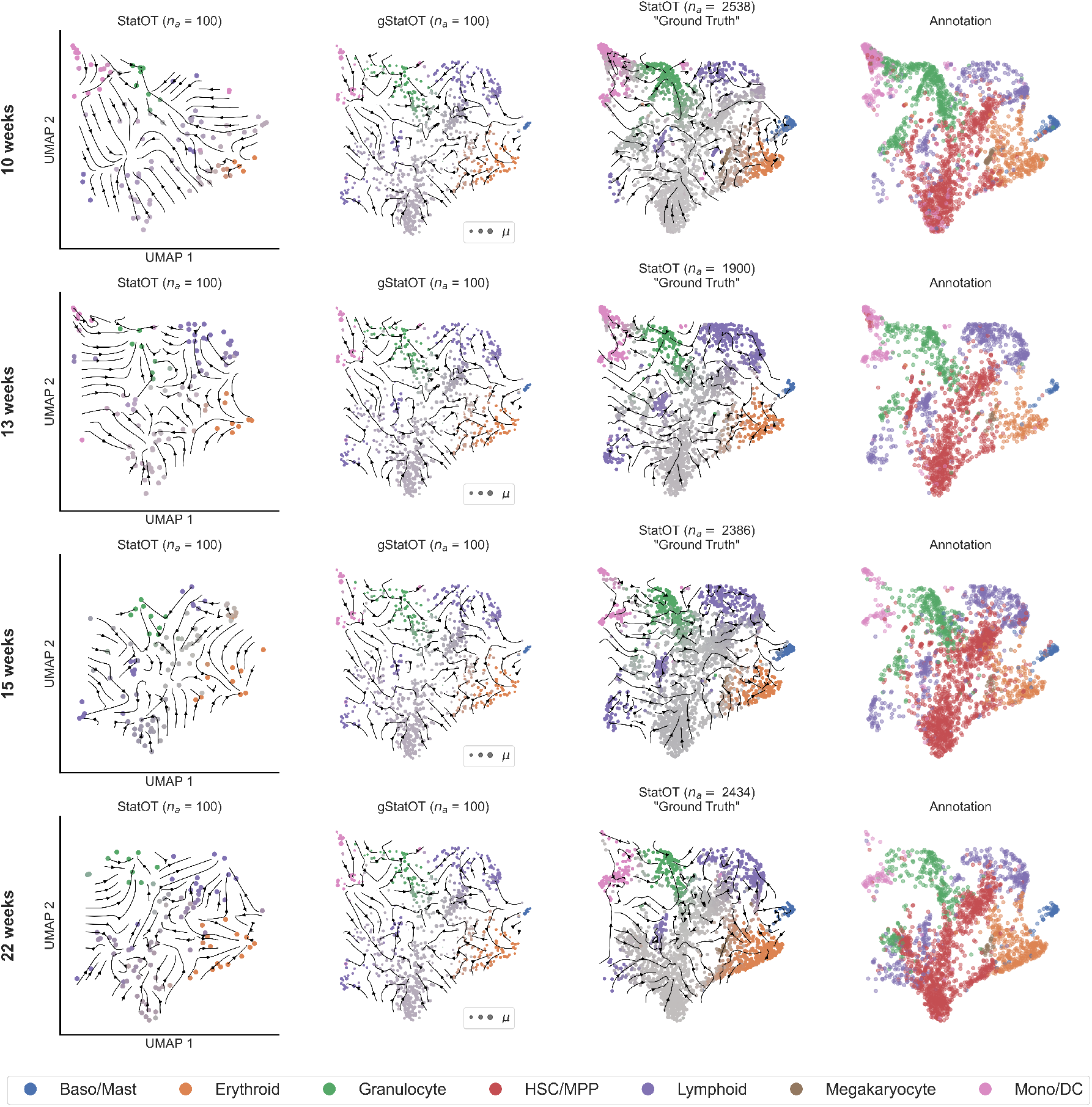
Same as Figure 7 **A**, but with the inferred flow fields and fate probabilities of gStatOT and StatOT at several prenatal ages.

**Figure SI.6:**
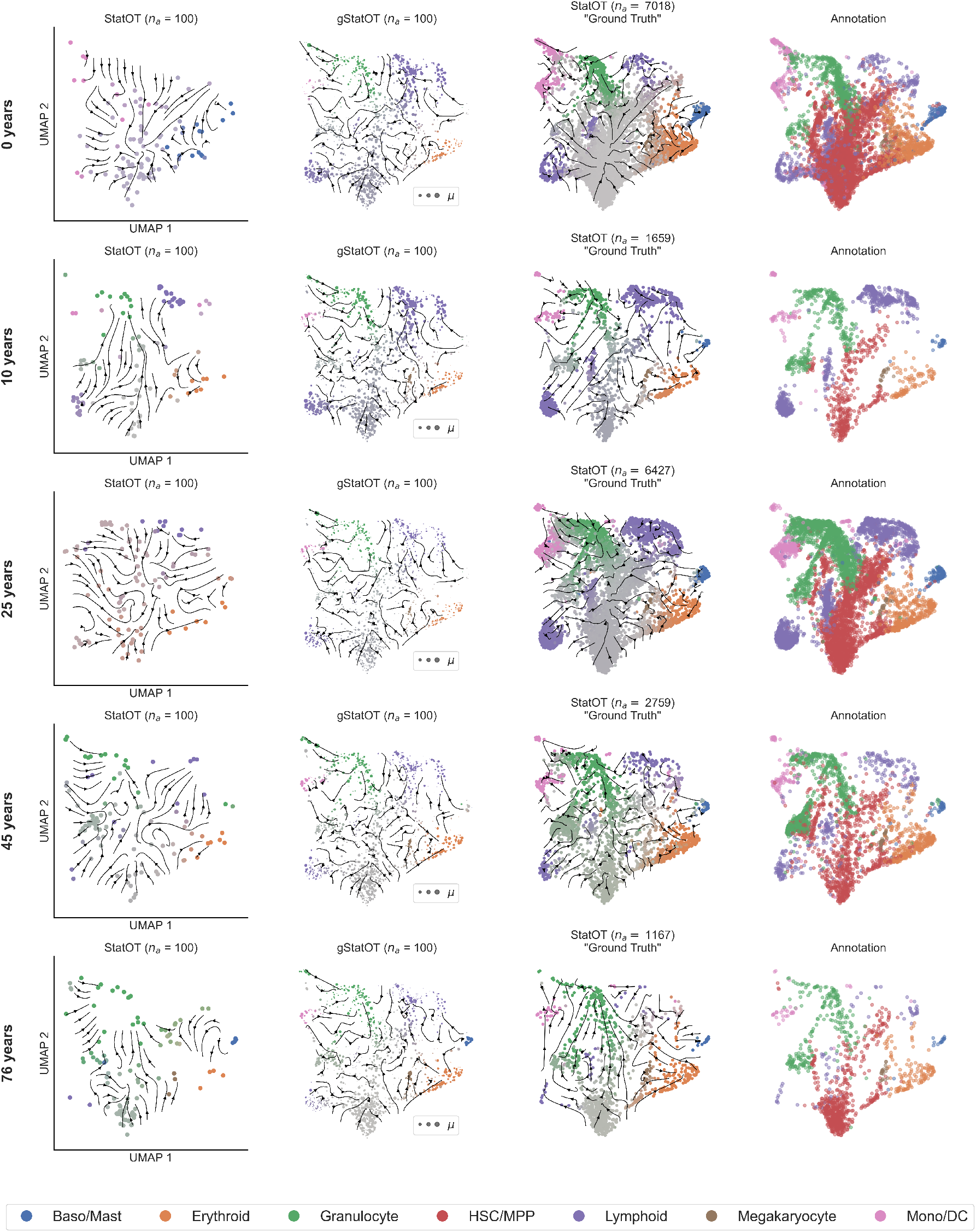
Same as Figure 7 **A**, but with the inferred flow fields and fate probabilities of gStatOT and StatOT at several postnatal ages.

**Figure SI.7:**
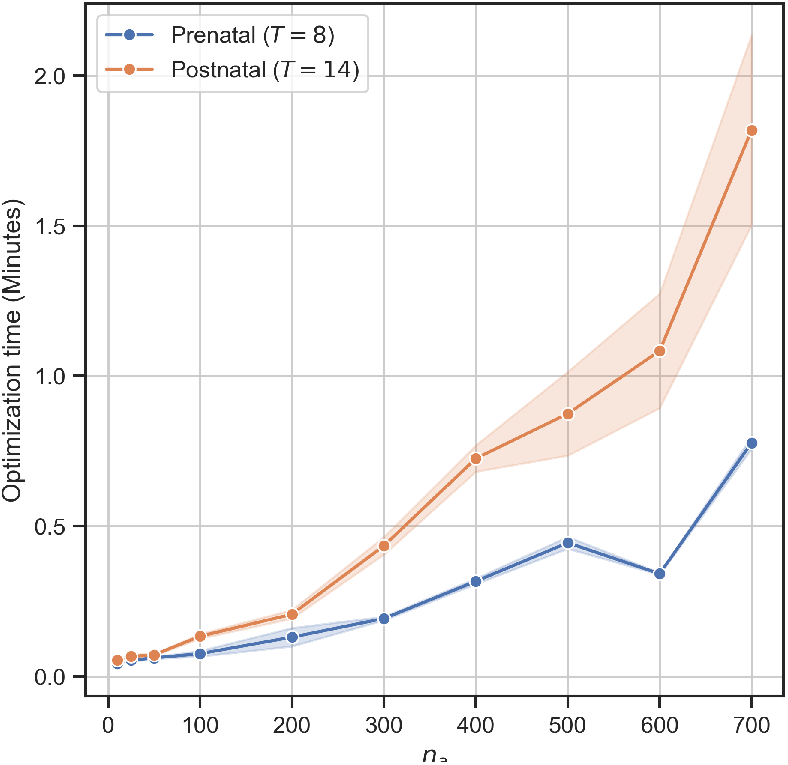
The time it took to solve gStatOT to produce the results in Figure 6 Panel **A** for each sparsity level, n_a_. The time was averaged over the 10 validation runs. Each run was performed on a single NVIDIA H100 80GB HBM3 MIG instance (profile: 3g.40gb).

## SI.6 Evaluation Metrics

### SI.6.1 Marginal Error

The second marginal of each optimal 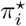 recovered by gStatOT provides an estimate of the population distribution at age *a*_*i*_. In order to test its debiasing ability, before applying the method to the simulated and real data, we downsample our initial samples and then compute the W_2_ distance between each 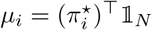 and the full sample of data, which we denote 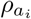. That is, we report

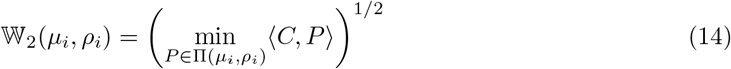

where *C*_*kl*_ = ∥*X*_*k*_ − *Y*_*l*_∥^2^, *X*_*k*_ ∈ supp *µ*_*i*_, *Y*_*l*_ ∈ supp *ρ*_*i*_. In Figure 4 panel **B** and Figure 6 panels **B, C**, we plot 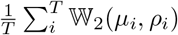 for each value of *T* .

### SI.6.2 Trajectory Metric

The Markov transition matrix *P*_*i*_ obtained from either StatOT, gStatOT, or PBA provides an estimate of the cell dynamics at age *a*_*i*_. We obtain a distribution, 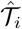, of cell trajectories at each age by sampling, 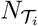, paths from *P*_*i*_ over a time *τ* starting from cells with positive growth. Letting 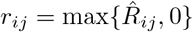, the initial cells at age *a*_*i*_ are sampled from the discrete distribution 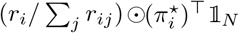. We report the W_2_ cost between 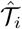 and the ground truth *T*_*i*_, which we describe obtaining in the text, with the cost

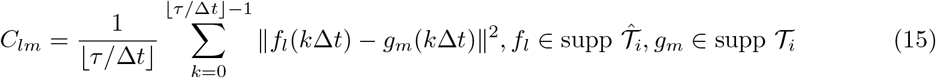

### SI.6.3 Fate probability metric

From each *P*_*i*_ we computed each cell’s lineage fate probabilities across *K* lineages, 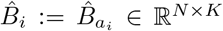, as in Section SI.3 for gStatOT/StatOT and used PBA’s built-in method to obtain its estimates. For each sampled cell, 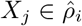, in the downsampled time series, their predicted lineage fate probability distribution, 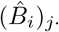, was compared to ground truth probabilities (*B*_*i*_)_*j*·_ using the total variation distance

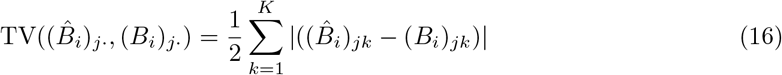

We report the average TV distance over all sampled cells at each age, 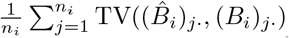 in Figures 4 and 5. In plots where we vary the number of time points, we report the average of this metric across all ages. As noted above, if a method failed to compute fate probabilities at a certain age, we set the total variation metric to its maximum value of 1 for that age.

### SI.6.4 Driver gene identification and average precision scoring

As done by Weiler et al. [S11], we use the correlation between gene expression and cell fate probability to produce ranked lists of putative driver genes at each age in the hematopoiesis time course. Before computing these correlations, we log-normalized the count data using the `normalize_total` (with `target_sum=`10^4^) and `log1p` functions from the `scanpy` Python package [S17].

Since gStatOT’s marginals are supported over all sampled data, we use a weighted correlation. Letting *G*_*l*_ be the column corresponding to a gene *l* in the *N* × *L* (log-normalized) gene expression matrix, we compute for weights *w* ∈ ℝ^*N*^ such that ∑_*j*_ *w*_*j*_ = 1

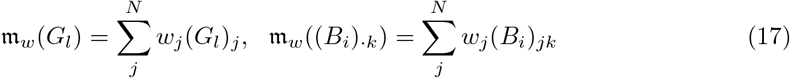

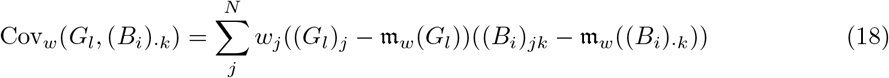

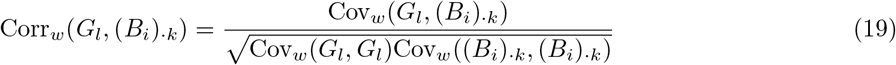

Recalling that 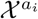 denotes the set of cells sampled at age *a*_*i*_, for StatOT we use the weights 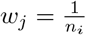 if cell 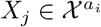 otherwise *w*_*j*_ = 0, while for gStatOT *w*_*j*_ = (*µ*_*i*_)_*j*_.

At each age, *a*_*i*_, and for each of the cell lineages we use our ground-truth estimates to form ranked lists, 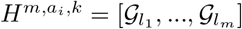, containing the top *m* = 10 genes most correlated with the *k*^th^ lineage. To evaluate a method’s ability to reconstruct these lists from downsampled data, we calculated the average precision (AP) [S18, S19] of each method’s list and reported the mean AP (mAP) over all ages in Figure 7 Panel **C**. In particular, given an estimated ranked list of driver genes for lineage *k*, 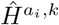, we evaluated

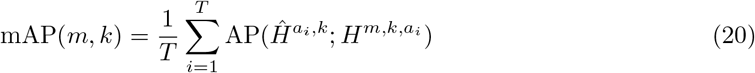

where for an estimated and ground truth lists *H* and *Ĥ*, respectively,

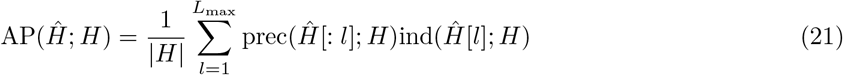

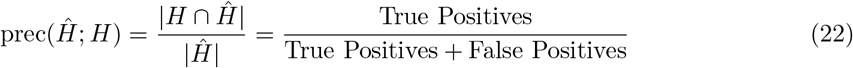

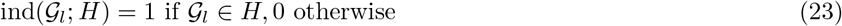

Here *H*[*l*] is the *l*^th^ element of *H*, and *H*[: *l*] denotes the first *l* items of the list. We set *L*_max_ = 1000, since relevant genes found outside this range will not significantly contribute to the average precision.

To provide a baseline measure of the ground-truth lists’ recoverability, we evaluated the temporal consistency (GT-TC) using the AP between the ground-truth lists at adjacent ages. Precisely,

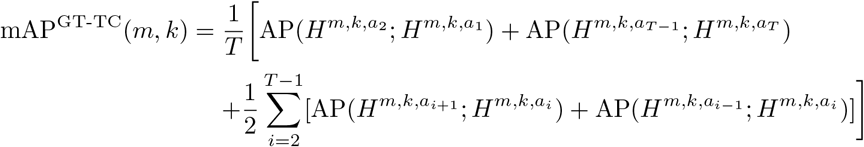

To compare with a simpler method, on the downsampled data, we obtained a ranked list of the top differentially expressed genes for each cell type relative to the rest of the data at each age using Scanpy’s [S17] `rank_genes_groups` function. We then computed the mAP using Equation 20.

## SI.7 Parameter sweeps

### SI.7.1 Synthetic data: Aging epigenetic landscape

**Figure SI.8:**
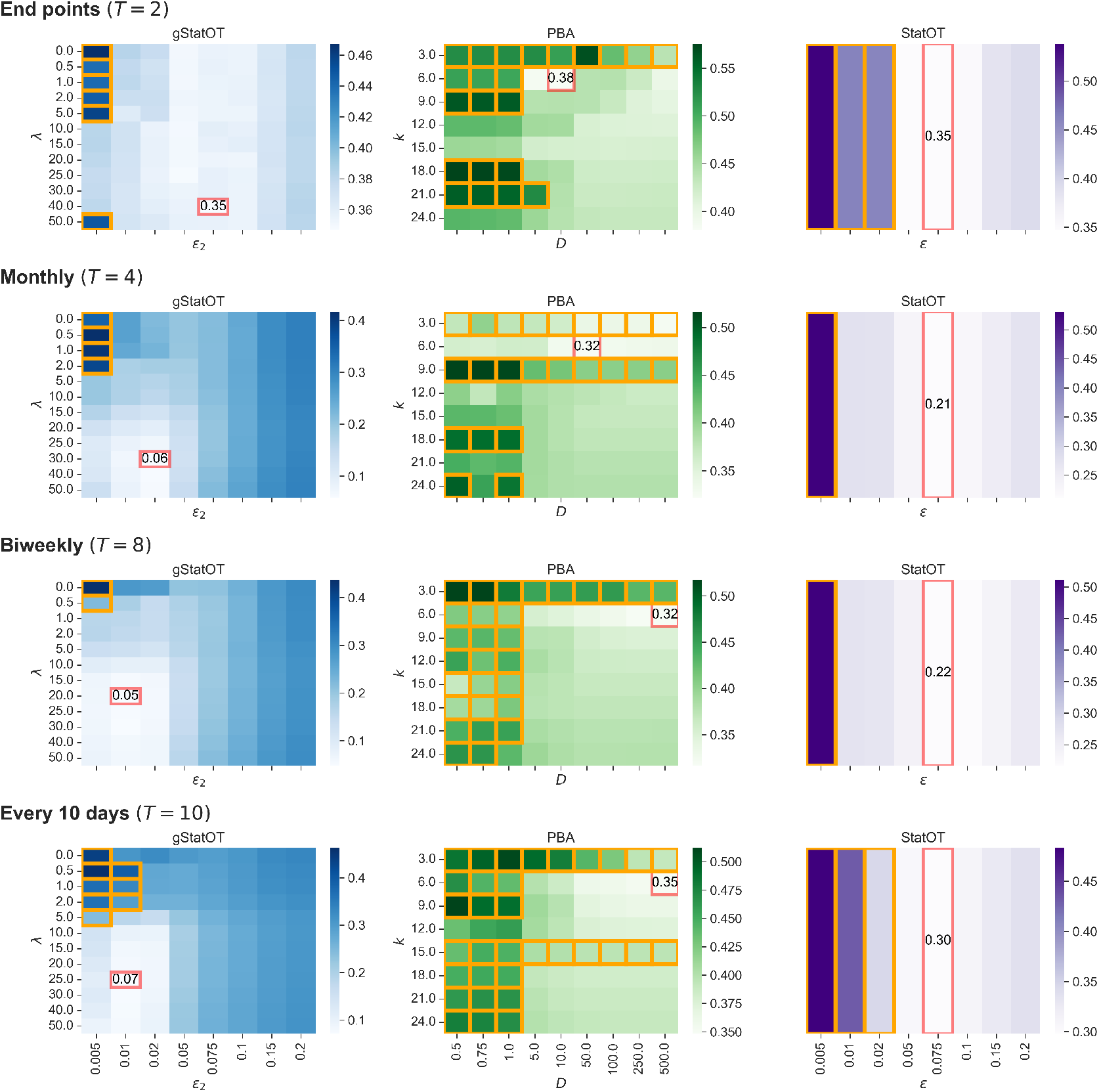
Average performance over the marginal, trajectory and fate probability metrics for parameter sweeps of gStatOT, StatOT, and PBA on our initial synthetic simulation. Parameters used for the results depicted in Figures 3 and 4 in the main text are highlighted in red. Parameter combinations that failed to compute fate probabilities at any age are highlighted in orange. Metrics were scaled by their minimum and maximum values over all method and parameter choices before taking the mean. Colour gradients are normalized for each method and value of T.

**Figure SI.9:**
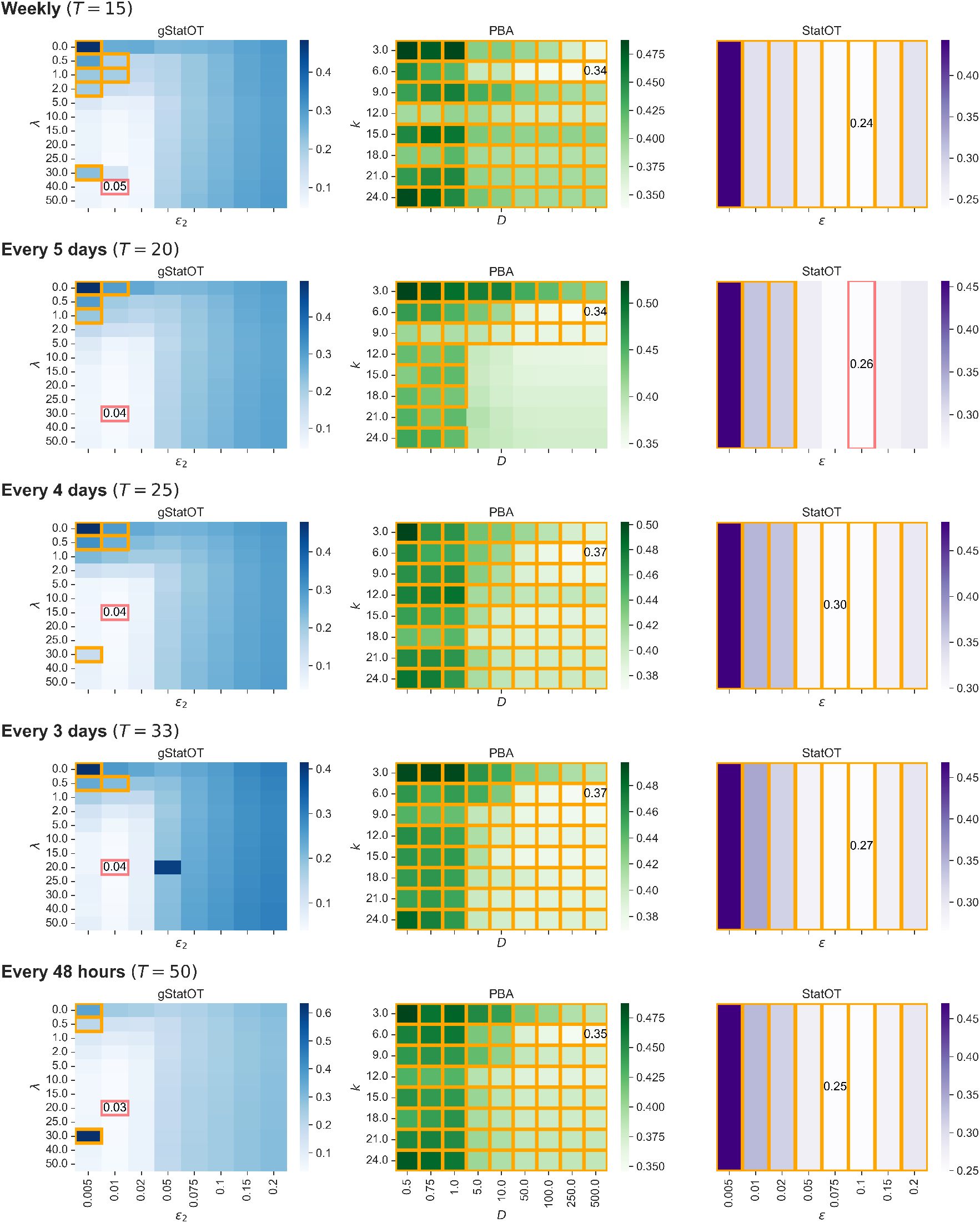
Continued from Figure SI.8, but for larger values of T.

### SI.7.2 Synthetic data: Loss of cellular identity over age induced by epigenetic erosion

**Figure SI.10:**
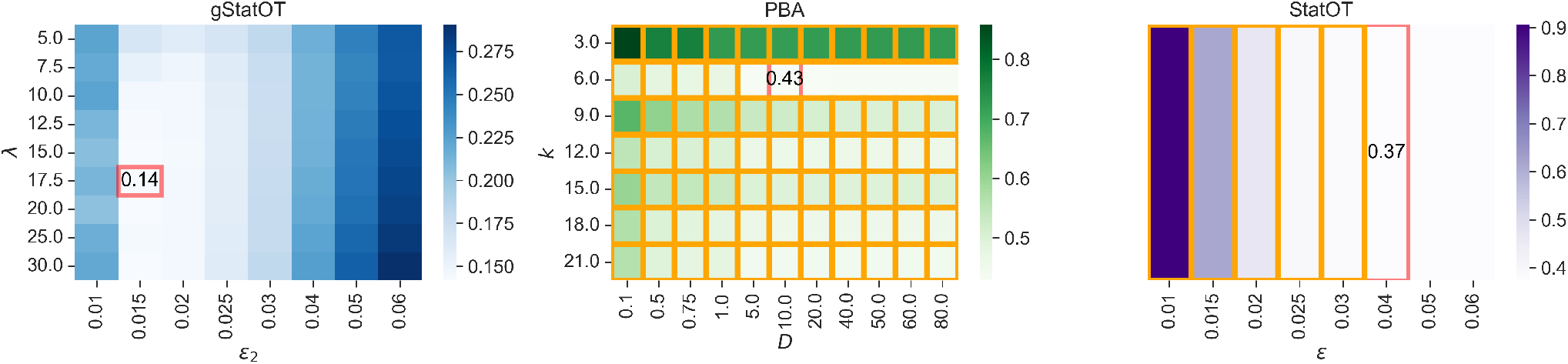
Fate probability TV error for parameter sweeps of gStatOT and StatOT on the simulated GRN data. As in the previous figures, the parameters used for the results depicted in Figure 5 in the main text are highlighted in red. Colour gradient is normalized over each method’s results.

### SI.7.3 Hematopoiesis

#### SI.7.3.1 Variable Sample Size

**Figure SI.11:**
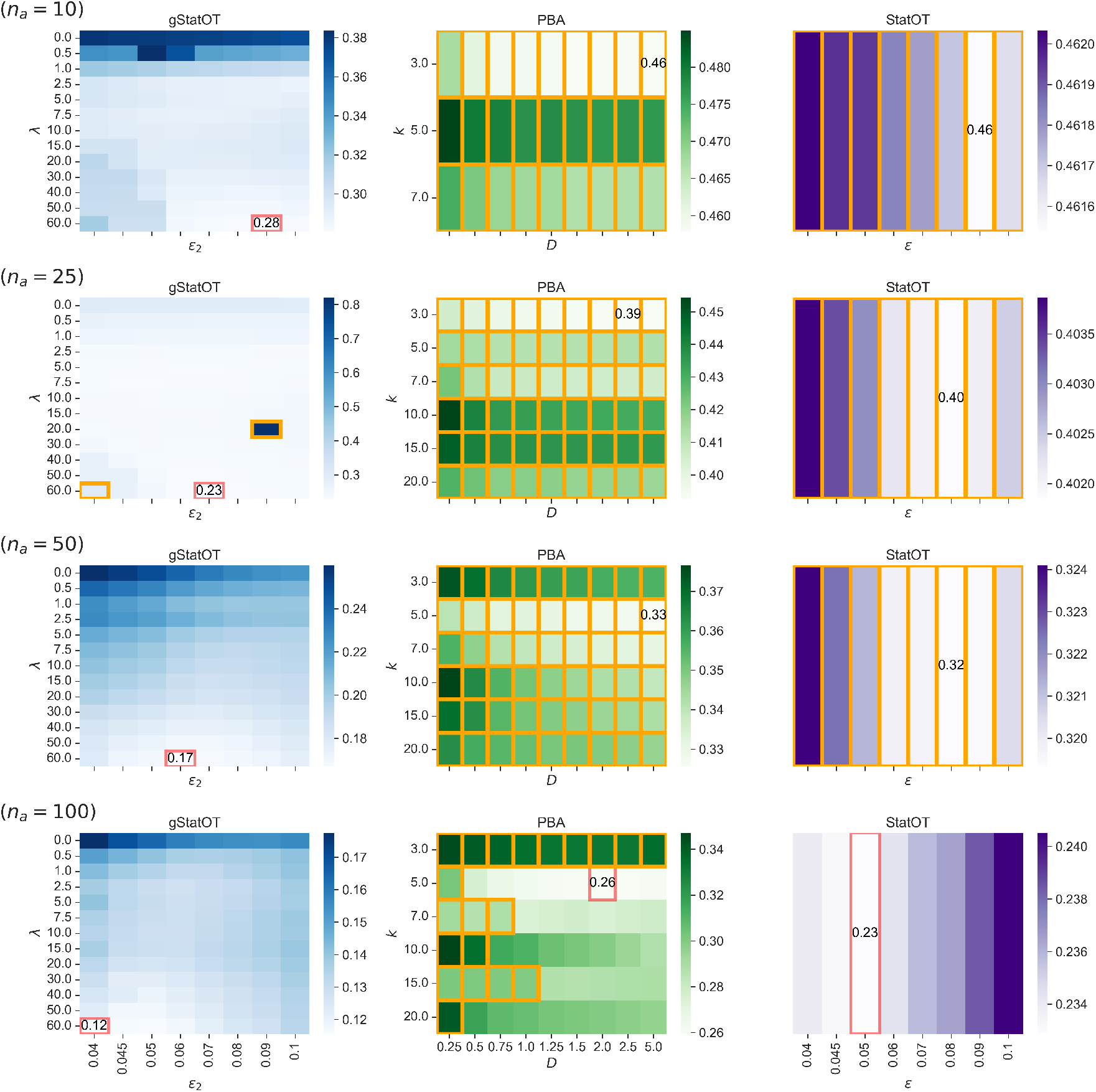
As in Figure SI.8, but for parameter sweeps of gStatOT, StatOT, and PBA on the down-sampled prenatal hematopoiesis data from Li et al. [S14], at varying sparsity levels. Metrics were scaled by their minimum and maximum values over all method and parameter choices before taking the weighted mean, valuing marginal reconstruction twice as much as the other metrics. The parameters used for the results displayed in Figures 6 and 7 in the text are highlighted in red. The colour gradient is normalized over gStatOT’s results for each value of n.

**Figure SI.12:**
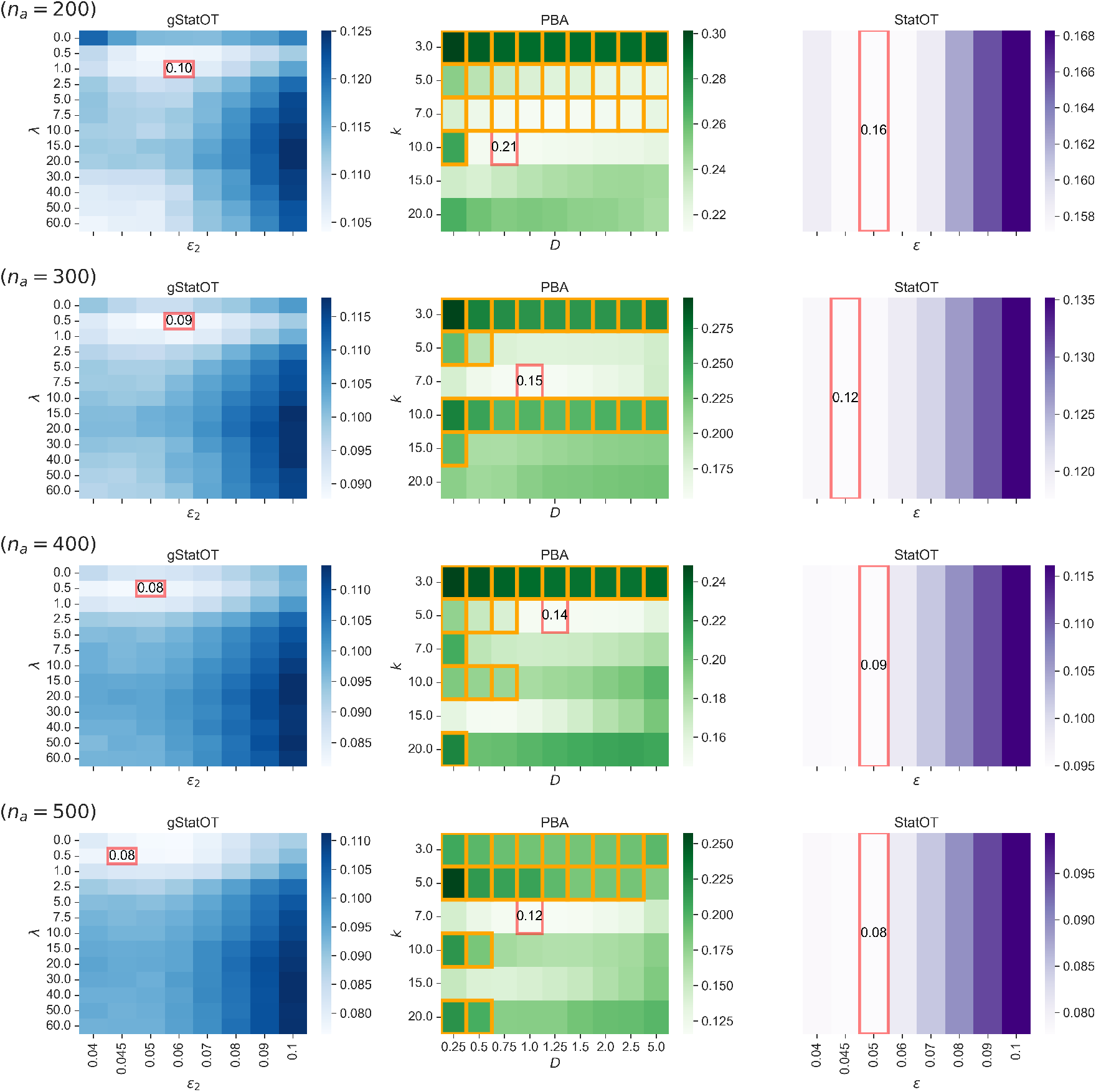
As in Figure SI.11, but for larger sample sizes.

**Figure SI.13:**
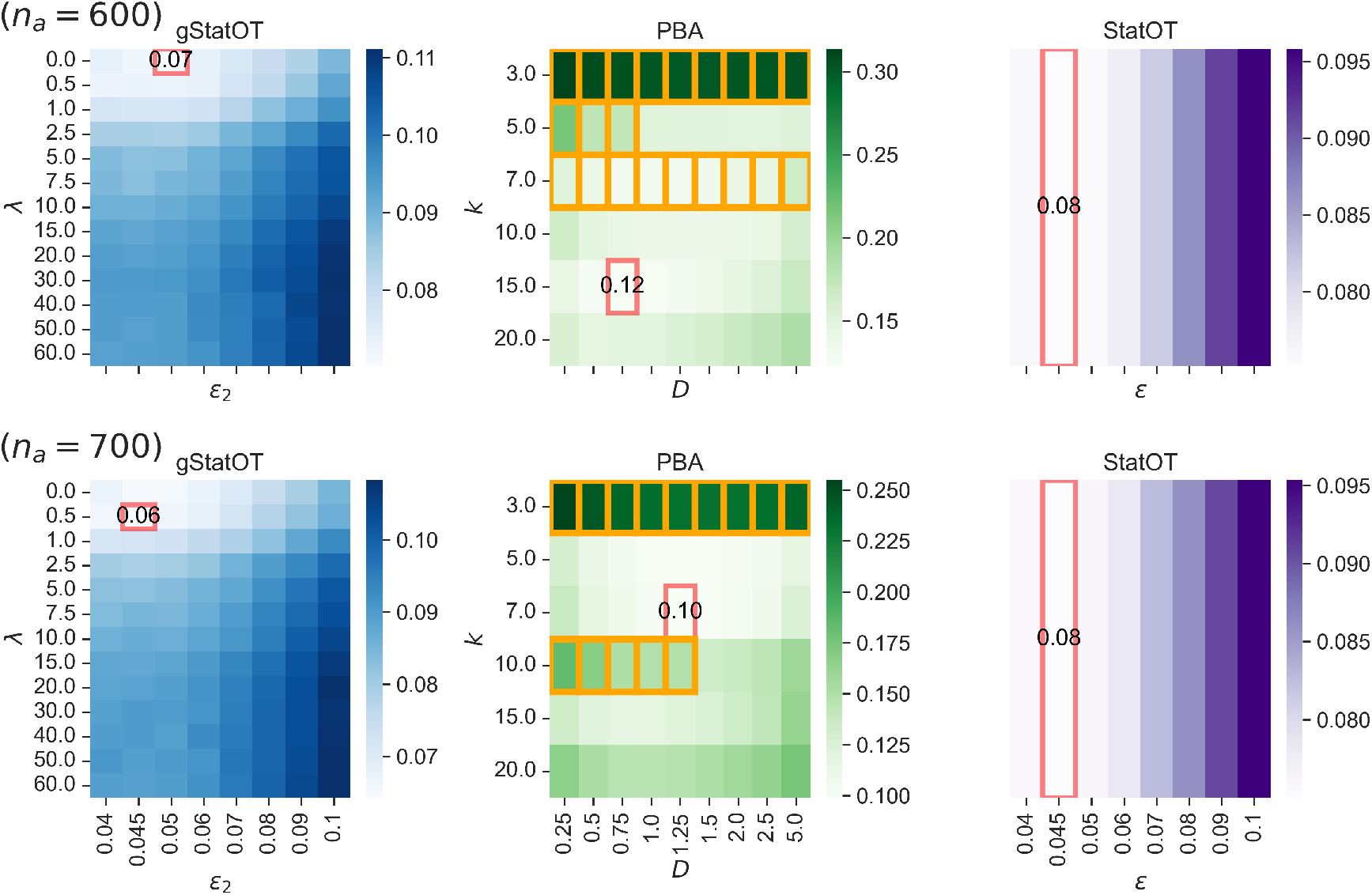
As in Figure SI.11, but for larger sample sizes.

**Figure SI.14:**
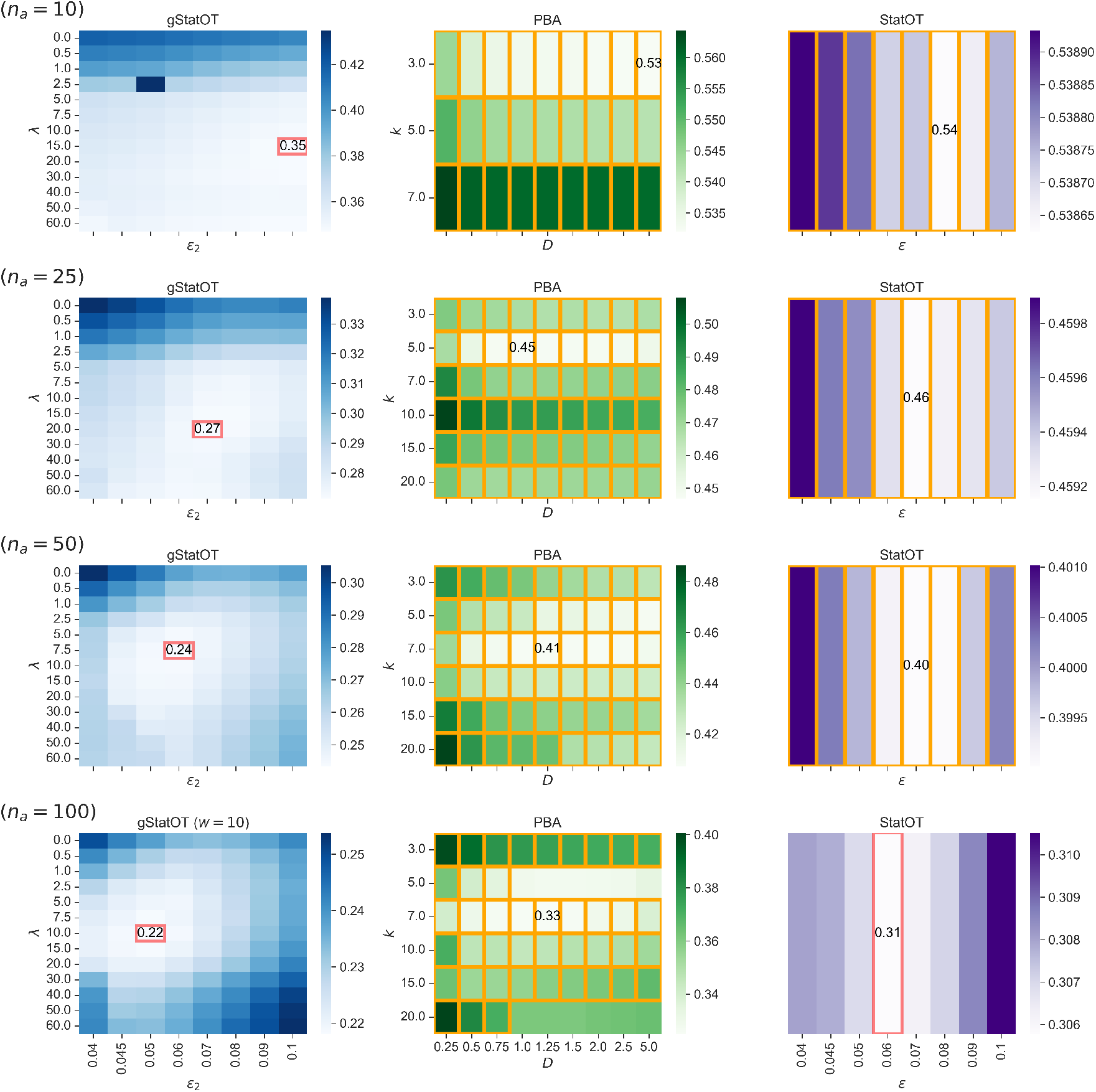
As in Figure SI.11, but for the postnatal time course.

**Figure SI.15:**
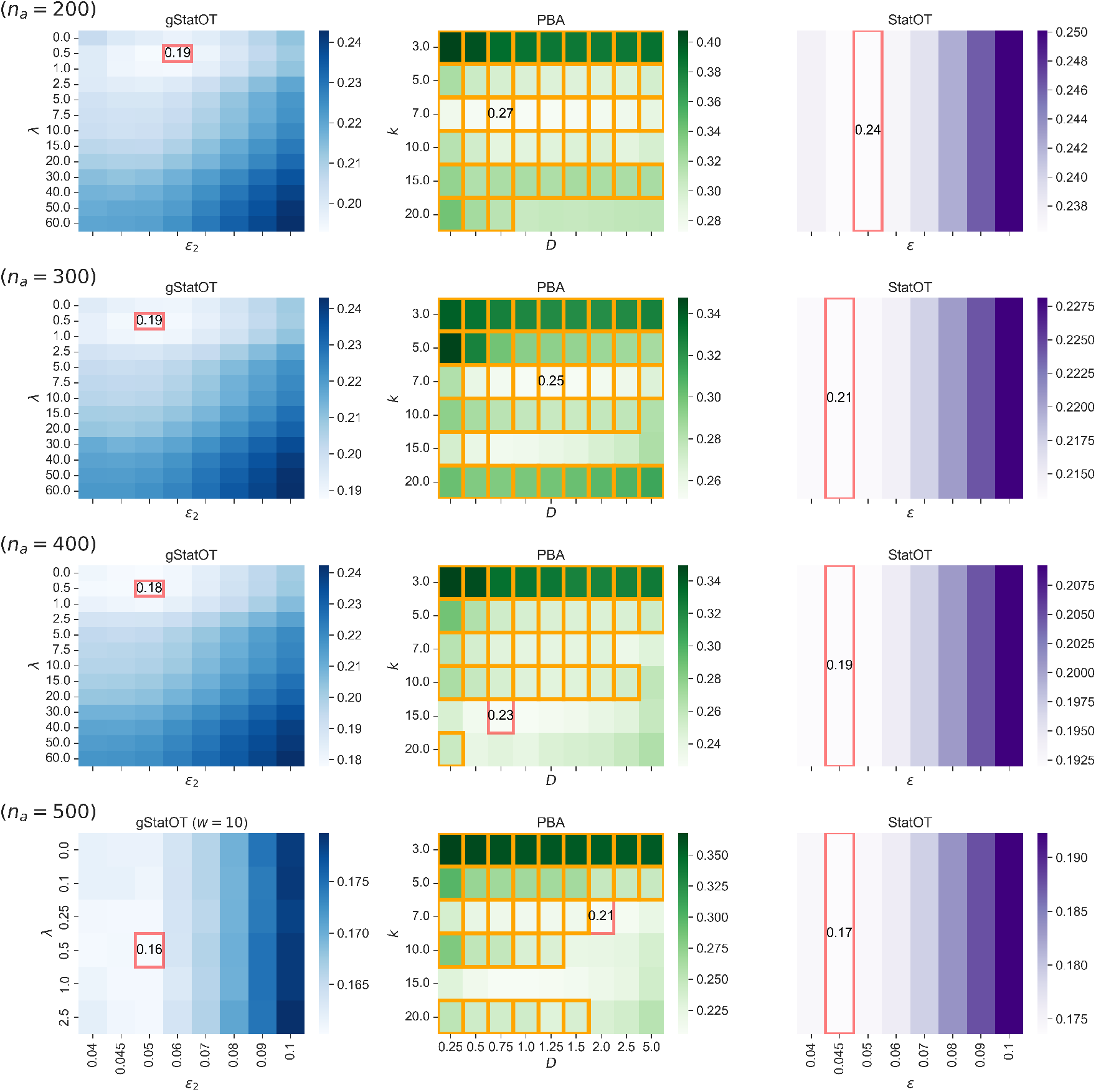
As in Figure SI.14, but for larger sample sizes. Note we have taken w = 10 when fitting gStatOT at n_a_ = 500 sparsity levels.

**Figure SI.16:**
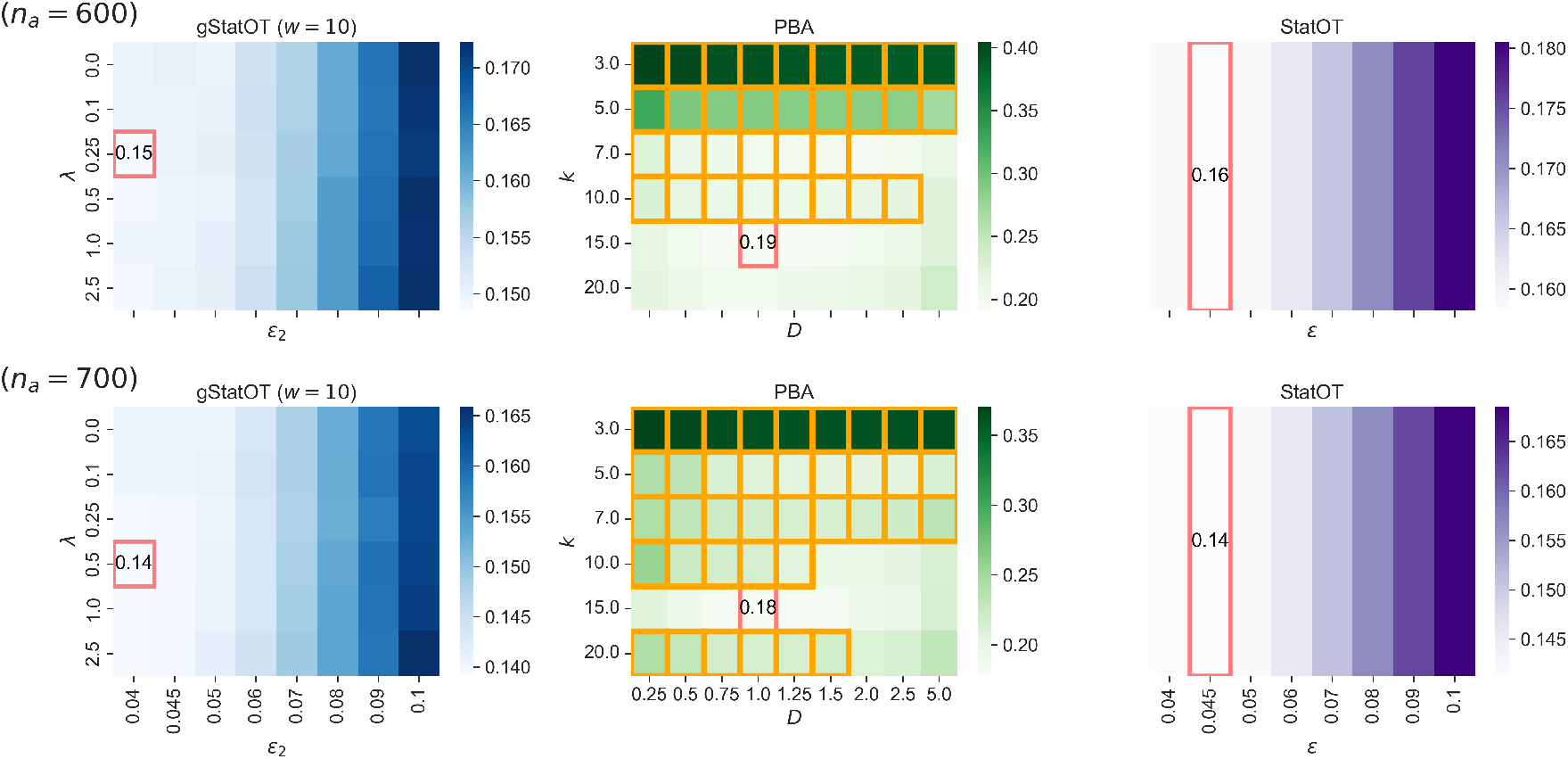
As in Figure SI.15, but for larger sample sizes. Note we have taken w = 10 when fitting gStatOT at these sparsity levels.

#### SI.7.3.2 Variable Number of Time Points

**Figure SI.17:**
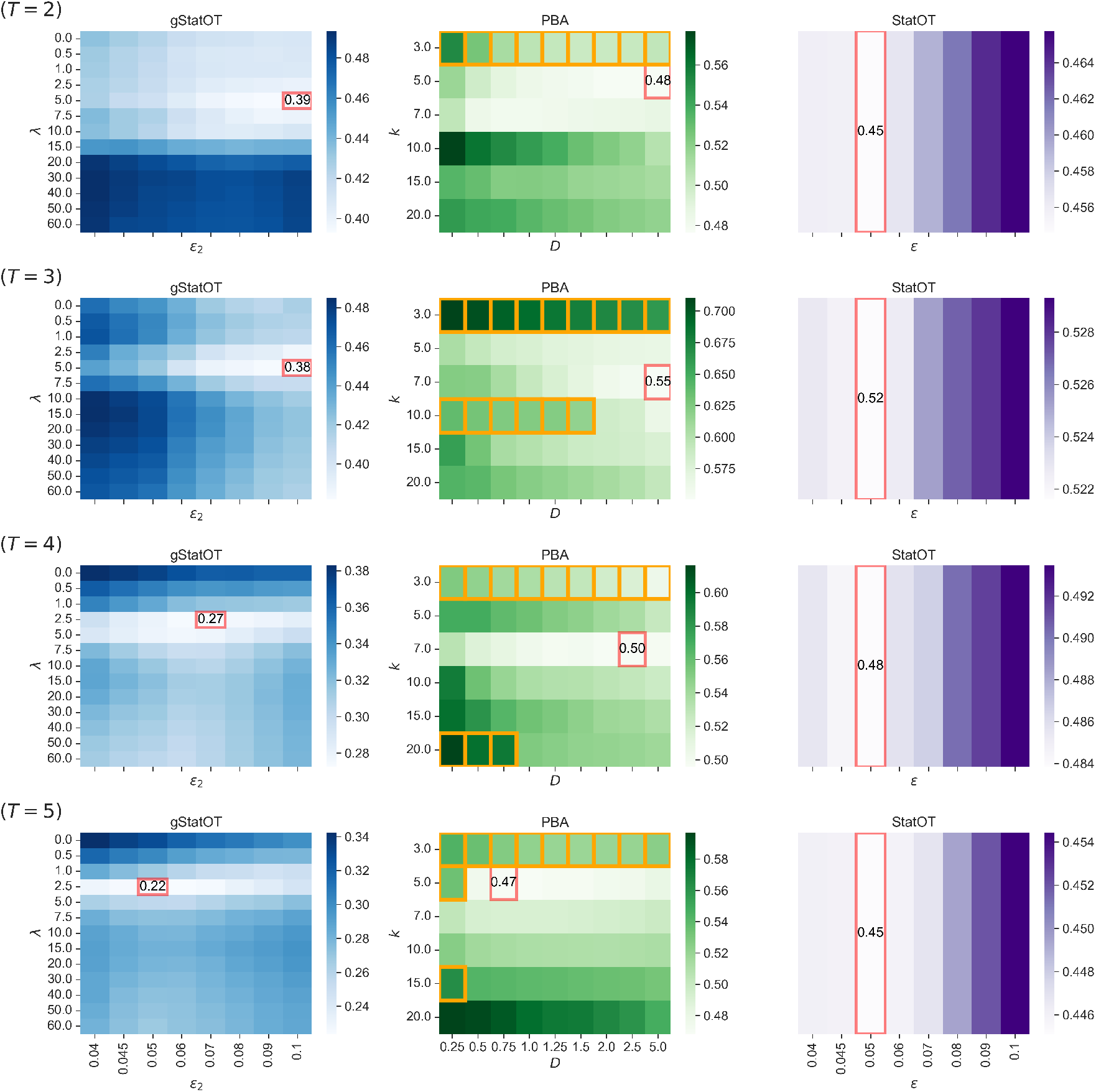
As in Figure SI.11, but for the prenatal time course with varying number of time points.

**Figure SI.18:**
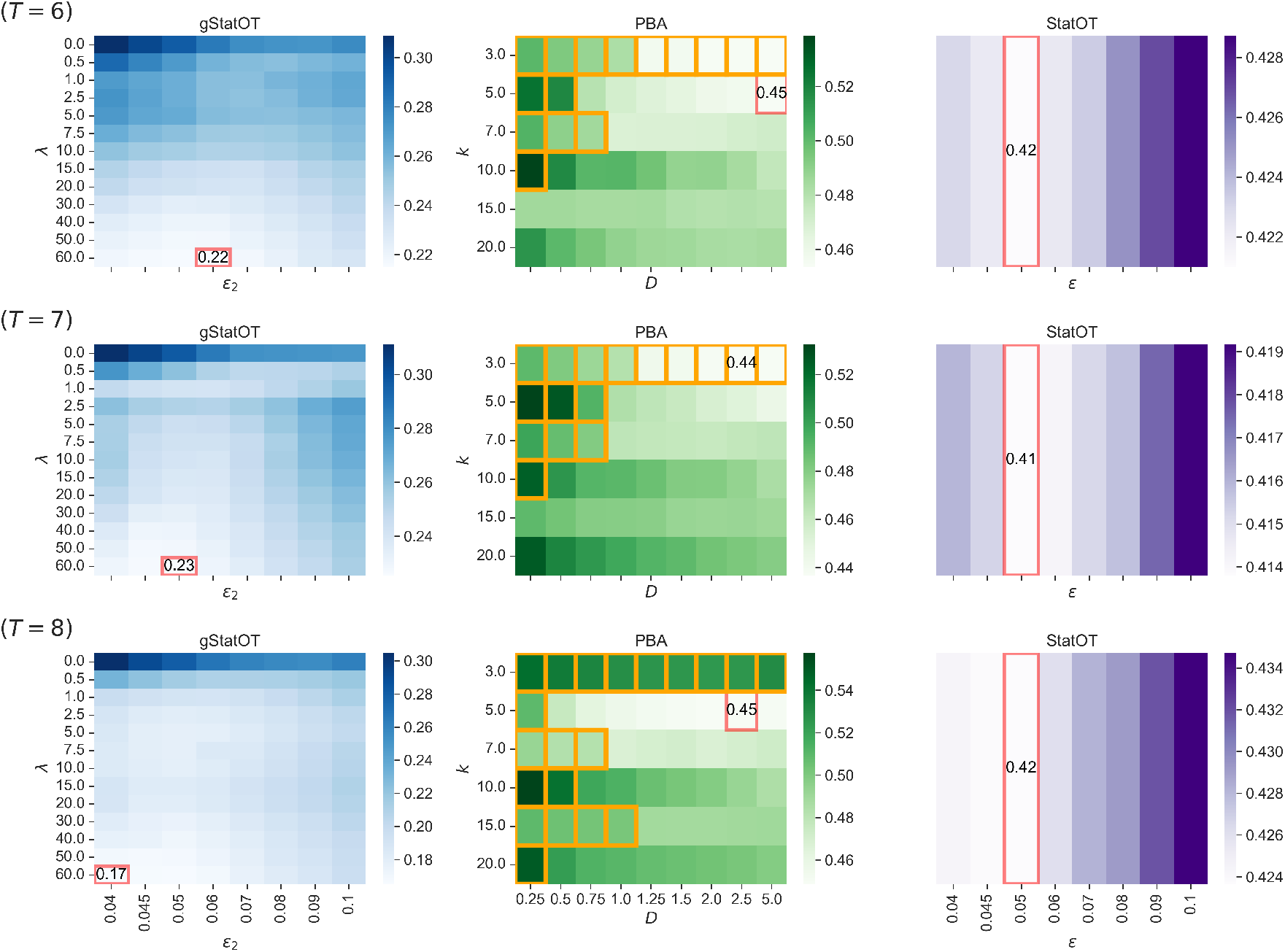
Continuation of Figure SI.17.

**Figure SI.19:**
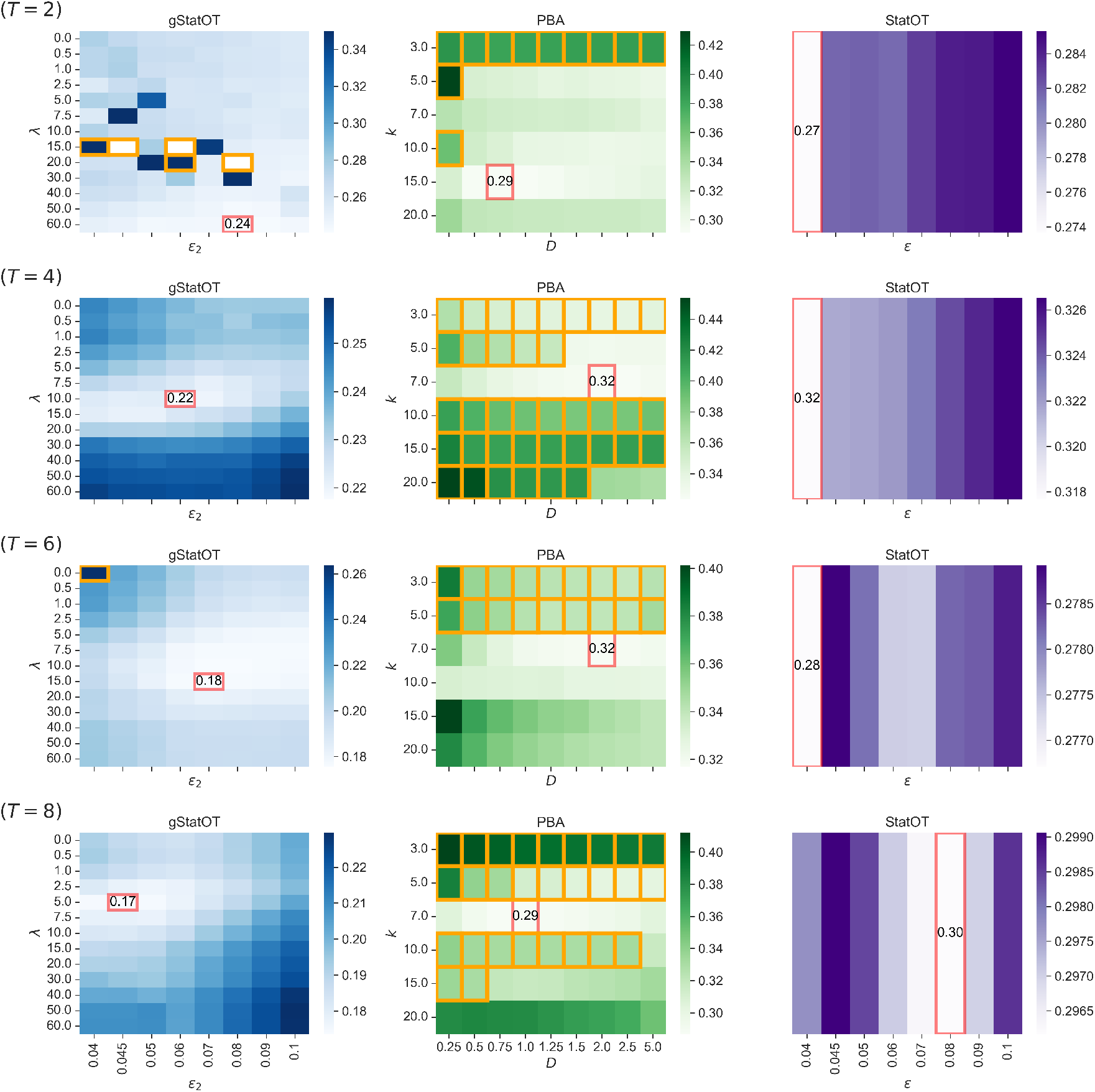
As in Figure SI.17, but for the postnatal time course.

**Figure SI.20:**
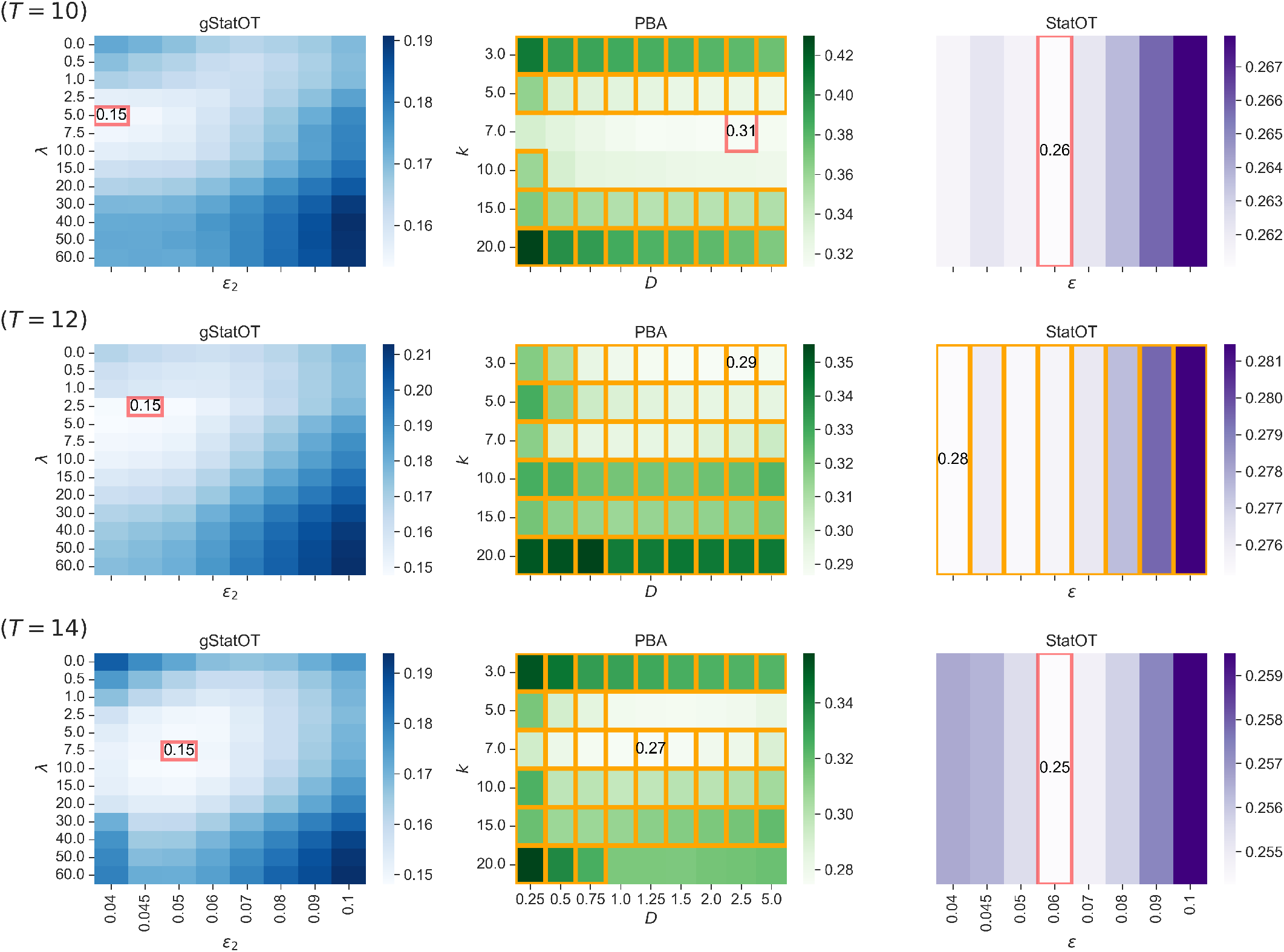
Continuation of Figure SI.19.

## SI.8 Usage of LLMs

Large Language models were utilized to assist in writing improved code and documentation at https://github.com/ColeBoyle/global-StationaryOT.

## Footnotes

∗ ∇_*z*_ denotes the gradient with respect to *z*

† We double the mass flow of sink states since they have naturally less out flow

## References

[1] Cole Trapnell, Davide Cacchiarelli, Jonna Grimsby, Prapti Pokharel, Shuqiang Li, Michael Morse, Niall J Lennon, Kenneth J Livak, Tarjei S Mikkelsen, and John L Rinn. The dynamics and regulators of cell fate decisions are revealed by pseudotemporal ordering of single cells. Nature biotechnology, 32(4):381–386, 2014.

[2] Kelly Street, Davide Risso, Russell B Fletcher, Diya Das, John Ngai, Nir Yosef, Elizabeth Purdom, and Sandrine Dudoit. Slingshot: cell lineage and pseudotime inference for single-cell transcriptomics. BMCcgenomics, 19(1):477, 2018.

[3] Volker Bergen, Marius Lange, Stefan Peidli, F Alexander Wolf, and Fabian J Theis. Generalizing rna velocity to transient cell states through dynamical modeling. Nature biotechnology, 38(12):1408–1414, 2020.

[4] Caleb Weinreb, Samuel Wolock, Betsabeh K Tusi, Merav Socolovsky, and Allon M Klein. Fundamental limits on dynamic inference from single-cell snapshots. Proceedings of the National Academy of Sciences, 115(10):E2467–E2476, 2018.

[5] Stephen Zhang, Anton Afanassiev, Laura Greenstreet, Tetsuya Matsumoto, and Geoffrey Schiebinger. Optimal transport analysis reveals trajectories in steady-state systems. PLoS computational biology, 17(12):e1009466, 2021.

[6] Philipp Weiler, Marius Lange, Michal Klein, Dana Pe’er, and Fabian Theis. Cellrank 2: unified fate mapping in multiview single-cell data. Nature Methods, 21(7):1196–1205, 2024.

[7] Hojun Li, Parker Côté, Michael Kuoch, Jideofor Ezike, Katie Frenis, Anton Afanassiev, Laura Greenstreet, Mayuri Tanaka-Yano, Giuseppe Tarantino, Stephen Zhang, et al. The dynamics of hematopoiesis over the human lifespan. Nature methods, 22(2):422–434, 2025.

[8] Alberto Patiño Douce. Thermodynamics of the Earth and Planets. Cambridge University Press, 2011.

[9] Johannes Hoppenau and Andreas Engel. On the work distribution in quasi-static processes. Journal of Statistical Mechanics: Theory and Experiment, 2013(06):P06004, 2013.

[10] Geoffrey Schiebinger, Jian Shu, Marcin Tabaka, Brian Cleary, Vidya Subramanian, Aryeh Solomon, Joshua Gould, Siyan Liu, Stacie Lin, Peter Berube, et al. Optimal-transport analysis of single-cell gene expression identifies developmental trajectories in reprogramming. Cell, 176(4):928–943, 2019.

[11] Alexander Tong, Jessie Huang, Guy Wolf, David Van Dijk, and Smita Krishnaswamy. Trajectorynet: A dynamic optimal transport network for modeling cellular dynamics. In International conference on machine learning, pages 9526–9536. PMLR, 2020.

[12] Hugo Lavenant, Stephen Zhang, Young-Heon Kim, Geoffrey Schiebinger, et al. Toward a mathematical theory of trajectory inference. The Annals of Applied Probability, 34(1A):428–500, 2024.

[13] Carlos López-Otín, Maria A Blasco, Linda Partridge, Manuel Serrano, and Guido Kroemer. The hallmarks of aging. Cell, 153(6):1194–1217, 2013.

[14] Zeming Wu, Jing Qu, and Guang-Hui Liu. Roles of chromatin and genome instability in cellular senescence and their relevance to ageing and related diseases. Nature Reviews Molecular Cell Biology, 25(12):979–1000, 2024.

[15] Bérénice A Benayoun, Elizabeth A Pollina, Param Priya Singh, Salah Mahmoudi, Itamar Harel, Kerriann M Casey, Ben W Dulken, Anshul Kundaje, and Anne Brunet. Remodeling of epigenome and transcriptome landscapes with aging in mice reveals widespread induction of inflammatory responses. Genome research, 29(4):697–709, 2019.

[16] Maria Luisa Amaral, Sainath Mamde, Michael Miller, Xiaomeng Hou, Jessica Arzavala, Julia Osteen, Nicholas D Johnson, Elizabeth Walker Smoot, Qian Yang, Emily Eisner, et al. Single-cell epigenomics uncovers heterochromatin instability and transcription factor dysfunction during mouse brain aging. Cell Reports, 45(3), 2026.

[17] Yuancheng Ryan Lu, Xiao Tian, and David A. Sinclair. The Information Theory of Aging. Nature Aging, 3(12):1486–1499, 2023.

[18] Yoshitsugu Oono and Marco Paniconi. Steady state thermodynamics. Progress of Theoretical Physics Supplement, 130:29–44, 1998.

[19] Luigi Ambrosio, Nicola Gigli, and Giuseppe Savaré. Gradient flows: in metric spaces and in the space of probability measures. Springer Science & Business Media, 2005.

[20] Ramin Rahni and Kenneth D Birnbaum. Week-long imaging of cell divisions in the arabidopsis root meristem. Plant methods, 15(1):30, 2019.

[21] Rachel Shahan, Che-Wei Hsu, Trevor M Nolan, Benjamin J Cole, Isaiah W Taylor, Laura Greenstreet, Stephen Zhang, Anton Afanassiev, Anna Hendrika Cornelia Vlot, Geoffrey Schiebinger, et al. A single-cell arabidopsis root atlas reveals developmental trajectories in wild-type and cell identity mutants. Developmental cell, 57(4):543–560, 2022.

[22] Helge Holden, Kenneth H Karlsen, Knut-Andreas Lie, and Nils Henrik Risebro. Splitting methods for partial differential equations with rough solutions. European Mathematical Society, 7:12, 2010.

[23] Christian Léonard. From the schrödinger problem to the monge–kantorovich problem. Journal of Functional Analysis, 262(4):1879–1920, 2012.

[24] Christian Léonard. A survey of the schrödinger problem and some of its connections with optimal transport. Discrete and Continuous Dynamical Systems, 34(4):1533–1574, 2014.

[25] Zhi-Quan Luo and Paul Tseng. On the convergence of the coordinate descent method for convex differentiable minimization. Journal of Optimization Theory and Applications, 72(1):7–35, 1992.

[26] James Bradbury, Roy Frostig, Peter Hawkins, Matthew James Johnson, Chris Leary, Dougal Maclaurin, George Necula, Adam Paszke, Jake VanderPlas, Skye Wanderman-Milne, and Qiao Zhang. JAX: composable transformations of Python+NumPy programs, 2025.

[27] Mathieu Blondel, Quentin Berthet, Marco Cuturi, Roy Frostig, Stephan Hoyer, Felipe Llinares-López, Fabian Pedregosa, and Jean-Philippe Vert. Efficient and modular implicit differentiation. Advances in neural information processing systems, 35:5230–5242, 2022.

[28] Marco Cuturi, Laetitia Meng-Papaxanthos, Yingtao Tian, Charlotte Bunne, Geoff Davis, and Olivier Teboul. Optimal transport tools (ott): A jax toolbox for all things wasserstein. arXiv preprint arXiv:2201.12324, 2022.

[29] Isaac Virshup, Sergei Rybakov, Fabian J Theis, Philipp Angerer, and F Alexander Wolf. anndata: Access and store annotated data matrices. Journal of Open Source Software, 9(101):4371, 2024.

[30] Larissa Traxler, Raffaella Lucciola, Joseph R. Herdy, Jeffrey R. Jones, Jerome Mertens, and Fred H. Gage. Neural cell state shifts and fate loss in ageing and age-related diseases. Nature Reviews Neurology, 19(7):434–443, 2023.

[31] Erin Connolly, Tony Pan, Maneesha Aluru, Sriram Chockalingam, Vishal Dhere, and Greg Gibson. Loss of immune cell identity with age inferred from large atlases of single cell transcriptomes. Aging Cell, 23(12):e14306, 2024.

[32] Tetsuya Ono and Richard G. Cutler. Age-dependent relaxation of gene repression: Increase of endogenous murine leukemia virus-related and globin-related RNA in brain and liver of mice. Proceedings of the National Academy of Sciences of the United States of America, 75(9):4431–4435, 1978.

[33] Philipp Oberdoerffer, Shaday Michan, Michael McVay, Raul Mostoslavsky, James Vann, SangKyu Park, Andrea Hartlerode, Judith Stegmuller, Angela Hafner, Patrick Loerch, et al. Sirt1 redistribution on chromatin promotes genomic stability but alters gene expression during aging. Cell, 135(5):907–918, 2008.

[34] Paul D Kaufman and Oliver J Rando. Chromatin as a potential carrier of heritable information. Current Opinion in Cell Biology, 22(3):284–290, 2010.

[35] Aditya Pratapa, Amogh P. Jalihal, Jeffrey N. Law, Aditya Bharadwaj, and T. M. Murali. Benchmarking algorithms for gene regulatory network inference from single-cell transcriptomic data. Nature Methods, 17(2):147–154, 2020.

[36] Sunnie Grace McCalla, Alireza Fotuhi Siahpirani, Jiaxin Li, Saptarshi Pyne, Matthew Stone, Viswesh Periyasamy, Junha Shin, and Sushmita Roy. Identifying strengths and weaknesses of methods for computational network inference from single-cell rna-seq data. G3: Genes, Genomes, Genetics, 13(3):jkad004, 2023.

[37] Carmen Bravo González-Blas, Seppe De Winter, Gert Hulselmans, Nikolai Hecker, Irina Matetovici, Valerie Christiaens, Suresh Poovathingal, Jasper Wouters, Sara Aibar, and Stein Aerts. Scenic+: single-cell multiomic inference of enhancers and gene regulatory networks. Nature methods, 20(9):1355–1367, 2023.

[38] Leland McInnes, John Healy, and James Melville. Umap: Uniform manifold approximation and projection for dimension reduction. arXiv preprint arXiv:1802.03426, 018.

[39] Kazuaki Kishida. Property of average precision and its generalization: An examination of evaluation indicator for information retrieval experiments. National Institute of Informatics Tokyo, Japan, 2005.

[40] Kevin P Murphy. Probabilistic machine learning: an introduction. MIT press, 2022.

[41] Dominik Klein, Giovanni Palla, Marius Lange, Michal Klein, Zoe Piran, Manuel Gander, Laetitia Meng-Papaxanthos, Michael Sterr, Lama Saber, Changying Jing, et al. Mapping cells through time and space with moscot. Nature, 638(8052):1065–1075, 2025.

[42] Seungbyn Baek and Insuk Lee. Single-cell atac sequencing analysis: from data preprocessing to hypothesis generation. Computational and structural biotechnology journal, 18:1429–1439, 2020.

## References

[S1] Hugo Lavenant et al. “Toward a mathematical theory of trajectory inference”. In: The Annals of Applied Probability 34.1A (2024), pp. 428–500.

[S2] Stephen P Boyd and Lieven Vandenberghe. Convex optimization. Cambridge university press, 2004.

[S3] Jonathan M Borwein and Qiji J Zhu. Techniques of variational analysis. Springer, 2005.

[S4] Gabriel Peyré, Marco Cuturi, et al. “Computational optimal transport: With applications to data science”. In: Foundations and Trends® in Machine Learning 11.5-6 (2019), pp. 355–607.

[S5] Zhi-Quan Luo and Paul Tseng. “On the convergence of the coordinate descent method for convex differentiable minimization”. In: Journal of Optimization Theory and Applications 72.1 (1992), pp. 7–35.

[S6] Stephen Wright, Jorge Nocedal, et al. “Numerical optimization”. In: Springer Science 35.67-68 (1999), p. 179.

[S7] James Bradbury et al. JAX: composable transformations of Python+NumPy programs. Version 0.7.1. 2025. URL: http://github.com/jax-ml/jax.

[S8] DeepMind et al. The DeepMind JAX Ecosystem. 2020. URL: http://github.com/google-deepmind.

[S9] Caleb Weinreb et al. “Fundamental limits on dynamic inference from single-cell snapshots”. In: Proceedings of the National Academy of Sciences 115.10 (2018), E2467–E2476.

[S10] Stephen Zhang et al. “Optimal transport analysis reveals trajectories in steady-state systems”. In: PLoS computational biology 17.12 (2021), e1009466.

[S11] Philipp Weiler et al. “CellRank 2: unified fate mapping in multiview single-cell data”. In: Nature Methods 21.7 (2024), pp. 1196–1205.

[S12] Marco Cuturi et al. “Optimal transport tools (ott): A jax toolbox for all things wasserstein”. In: arXiv preprint arXiv:2201.12324 (2022).

[S13] Aditya Pratapa et al. “Benchmarking Algorithms for Gene Regulatory Network Inference from Single-Cell Transcriptomic Data”. In: Nature Methods 17.2 (2020), pp. 147–154. ISSN: 1548-7091, 1548-7105. doi: 10.1038/s41592-019-0690-6.

[S14] Hojun Li et al. “The dynamics of hematopoiesis over the human lifespan”. In: Nature methods 22.2 (2025), pp. 422–434.

[S15] Yuhan Hao et al. “Dictionary learning for integrative, multimodal and scalable single-cell analysis”. In: Nature Biotechnology (2023). doi: 10.1038/s41587-023-01767-y. URL: https://doi.org/10.1038/s41587-023-01767-y.

[S16] J. D. Hunter. “Matplotlib: A 2D graphics environment”. In: Computing in Science & Engineering 9.3 (2007), pp. 90–95. doi: 10.1109/MCSE.2007.55.

[S17] F Alexander Wolf, Philipp Angerer, and Fabian J Theis. “SCANPY: large-scale single-cell gene expression data analysis”. In: Genome biology 19.1 (2018), p. 15.

[S18] Kazuaki Kishida. Property of average precision and its generalization: An examination of evaluation indicator for information retrieval experiments. National Institute of Informatics Tokyo, Japan, 2005.

[S19] Kevin P Murphy. Probabilistic machine learning: an introduction. MIT press, 2022.

